# Development of EphA2 siRNA-loaded lipid nanoparticles and combination with a small-molecule histone demethylase inhibitor in prostate cancer cells and tumor spheroids

**DOI:** 10.1101/2020.09.28.315341

**Authors:** Ezgi Oner, Mustafa Kotmakci, Anne-Marie Baird, Steven G. Gray, Bilge Debelec Butuner, Emir Bozkurt, Ayse Gulten Kantarci, Stephen P. Finn

## Abstract

siRNAs hold a great potential for cancer therapy, however, poor stability in body fluids and low cellular uptake limit their use in the clinic. To enhance the bioavailability of siRNAs in tumors, novel, safe, and effective carriers are needed. Here, we developed cationic solid lipid nanoparticles (cSLNs) to carry siRNAs targeting EphA2 receptor tyrosine kinase (siEphA2), which is overexpressed in many solid tumors including prostate cancer (PCa). Using DDAB cationic lipid instead of DOTMA reduced nanoparticle size and enhanced both cellular uptake and gene silencing in PCa cells. After verifying the efficacy of siEphA2-loaded nanoparticles, we further evaluated a potential combination with a histone lysine demethylase inhibitor, JIB-04. Silencing EphA2 by siEphA2-loaded DDAB-cSLN did not affect the viability (2D and 3D), migration, and clonogenicity of PC-3 cells alone. However, upon co-administration, there was a decrease in the aforementioned cellular responses due to JIB-04. Furthermore, JIB-04 decreased EphA2 expression, and thus, silencing efficiency of siEphA2-loaded nanoparticles was further increased with co-treatment. In conclusion, we have successfully developed a novel siRNA-loaded lipid nanoparticle for targeting EphA2. Moreover, detailed preliminary results of the effects of JIB-04, alone and in combination with siEphA2, on PCa cells and tumor spheroids were presented for the first time. Our delivery system provides high transfection efficiency and shows a great promise for targeting other genes and cancer types in further *in vitro* and *in vivo* studies.

## 1. Introduction

Cancer is a leading cause of death worldwide, with prostate cancer accounting for 7.1% of all cancer cases [1]. Today, traditional chemotherapeutic agents commonly used in treatment have serious side effects due to their toxicity on healthy cells [2]. Thus, new agents with high selectivity are being investigated as better substitutes for chemotherapeutics to improve outcomes and quality of life. Because cancer has a complex heterogeneity, recent studies have focused on personalized medicine strategies which specifically target tumor markers [3,4]. These innovative anticancer treatment approaches include the use of RNA interference (RNAi)-based therapeutics that specifically silence oncogenes [5] or the use of selective anticancer agents that reorganize the dysregulated epigenetic mechanism in tumor cells [6].

Small interfering RNA (siRNA) is an effective tool of RNAi-based therapy for gene silencing with high specificity and efficacy [7]. However, siRNA molecules require suitable carriers (viral vectors or non-viral delivery systems) to be successfully translated into the clinic. These carriers are essential for the stability of siRNA molecules in body fluids, efficient uptake, and subsequent intracellular trafficking for desired efficacy [8]. Among these delivery systems, lipid-based non-viral carriers have attracted great attention as they provide a safe alternative to viral vectors, and patisiran (ONPATTRO^®^, formerly ALN-TTR02), the first-in-human lipid nanoparticle-based siRNA drug, received US-FDA approval in 2018 [9]. Additionally, many clinical trials are ongoing with lipid-based siRNA carriers for the treatment of various diseases including prostate cancer [10,11]. One of those being tested is an siRNA targeting EphA2/ 1,2-Dioleoyl-sn-glycero-3-phosphocholine (siEphA2/ DOPC) liposome, EPHARNA, which silences high levels of EphA2 receptor in advanced or recurrent solid tumors including prostate [12]. Cationic solid lipid nanoparticles (cSLNs) have many advantages over liposomal carriers such as physical stability, preparation without organic solvents, cost-effectiveness, and ease of scale-up [13]. However, to the best of our knowledge, there is no study using cSLNs for siEphA2 delivery.

Eph receptor A2 (EphA2) is a member of the receptor tyrosine kinase (RTK) family that regulates key cellular processes such as cell proliferation, survival, and differentiation [14]. EphA2 is overexpressed in many cancers such as prostate, ovarian, breast, lung, brain, urothelial, and skin [15]. EphA2 upregulation is associated with tumor invasion, metastasis, survival, and angiogenesis [14,15]. Therefore, various systems targeting EphA2 and other RTKs have been developed for the treatment of many solid tumors including prostate cancer [15,16]. However, resistance to RTK inhibitors can occur in tumors through many different mechanisms. One mechanism involves the rewiring of epigenetic regulation in cancer cells. For instance, overexpression of the histone lysine demethylase (KDM)5A is indispensable for the emergence of a subpopulation of cancer cells resistant to EGFR inhibitors [17]. In addition, other members of the KDM family (KDM4A, KDM4B, and KDM4C) are overexpressed in tumors of prostate, lung, colorectal, breast [18] and have been shown to promote drug resistance [19]. As such, various KDM inhibitors have been developed to reverse the epigenetic rewiring [20] and holds a great potential for single treatment as well as combination therapy [21,22]. JIB-04 (5-chloro-N-[(E)-[phenyl(pyridin-2-yl)methylidene]amino]pyridin-2-amine) is a novel small-molecule inhibitor of KDM5A-B, KDM4A-E, KDM6B that shows selectivity to cancer cells without harming normal cells [23,24]. The anti-tumor effects of JIB-04 in lung [22,23,25], glioblastoma [21,26], colorectal [27], gastric [28] cancers, hepatocellular carcinoma [29], leukemia [30], and Ewing sarcoma [31] have been studied in a great detail. JIB-04 re-sensitizes resistant cells to carboplatin-paclitaxel, cytarabine, and temozolomide; and increases the anti-tumor activity of these drugs in non-small cell lung cancer (NSCLC) [22], leukemia [30], and glioblastoma multiforme [21,26], respectively. However, studies with JIB-04 in prostate cancer are only limited to *in vitro* cytotoxicity [23].

In this study, we aimed to develop a cationic solid lipid nanoparticle/siRNA complex targeting EphA2 receptor (cSLN/siEphA2 complex) for use in the treatment of advanced cancers with high levels of EphA2 expression and investigate its anticancer activity alone and in combination with pan-KDM inhibitor JIB-04 for prostate cancer therapy *in vitro*. After characterizing and selecting the most effective complex based on cellular uptake efficiency, cytotoxicity, and EphA2 gene silencing efficiency in prostate cancer cells, its anticancer effect with epigenetic agent JIB-04 on cell viability (2D and 3D tumor spheroids), migration, and colony formation was evaluated *in vitro*.

## 2. Materials and Methods

### 2.1. Materials

JIB-04 was a generous gift from Assoc. Prof. PhD Elisabeth Martinez from Department of Pharmacology, University of Texas Southwestern Medical Center, USA. Solid lipids – Precirol ATO 5 and Compritol 888 ATO were gifts from Gattefossé (France). Surfactants – Kolliphor RH 40 (Cremophor RH 40, KRH40) was a gift from BASF (Germany) and Span 80 (S80) was purchased from Sigma-Aldrich (Germany). Propylene glycol (PG) was used as a co-surfactant and acquired from Merck (Germany). Cationic lipids – 1,2-di-O-octadecenyl-3-trimethylammonium propane (chloride salt) (DOTMA) was purchased from Avanti Polar Lipids (USA) and dimethyldioctadecylammonium bromide (DDAB) from Tokyo Chemical Industry (Japan). Standard ultra-pure water (upH_2_O) was used in all formulations.

### 2.2. Preparation of cSLNs by a modified hot microemulsion method

Formulations were prepared by the modified hot microemulsion method as described previously [32]. Briefly, solid lipids (Precirol: Compritol; 1.25%: 1.25%; w/w), the mixture of surfactants and co-surfactant (KRH40: S80: PG; 1.5%: 0.5%: 2%; w/w), and cationic lipid DDAB for DDAB-cSLN or DOTMA for DOTMA-cSLN (0.5% w/w) were weighed into a glass vial and stirred at the temperature above the melting points of solid lipids (80°C). After obtaining a homogenous mixture, upH_2_O at 80°C was added dropwise to this mixture while stirring (1500 rpm). Subsequently, the sample was cooled by stirring for an extra 30 min at room temperature.

### 2.3. Measurement of particle size and zeta potential

The cSLN formulations and their siRNA complexes were diluted in upH_2_O as appropriate. The dynamic light scattering (DLS) method was performed at 173° back-scattering mode to measure the Z-average particle size and polydispersity index (PDI) using Zetasizer NanoZS instrument (Malvern Panalytical Ltd., UK). Electrophoretic light scattering technique was used to measure electrophoretic mobility and zeta potential was calculated according to the Smoluchowski equation using equipment software (Malvern Panalytical Ltd., UK). To determine the change in particle size, PDI, and zeta potential of cSLN formulations upon storage at room temperature, they were stored in 4 mL clear glass vials with polytetrafluoroethylene-lined caps (Agilent, USA), and measurements were performed at 2, 4, 8 and 12 weeks after the preparation day (Day 0). All measurements were performed at 25°C at least in triplicate.

### 2.4. Gel retardation (complexation) assay

Gel retardation (shift) assay was conducted using agarose gel electrophoresis to determine the optimal amount of the formulation for complex formation with siRNA [32]. Briefly, different amounts of cSLN with a constant amount of siRNA (67 ng) were incubated at room temperature on a shaker (500 rpm, 30 min). Complex formation was evaluated by observing the electrophoretic mobility of free siRNA in 2% agarose gel. The gel was visualized under a UV Transilluminator (Vilber Lourmat, France) after ethidium bromide (0.5 µg/mL) staining. The molar ratio of cationic lipid nitrogens to siRNA phosphates (N/P ratio) with no detectable band or smear of free siRNAs was chosen as the optimum and used in further analyses.

### 2.5. RNase A and serum protection assay

The pre-formed cSLN/siRNA complexes were incubated at 37°C with RNase A (10 µg/mL) for 30 min or with fetal bovine serum (FBS; #10270-106 Gibco, 50% v/v) for 1 h or 4 h. After incubation, a stop solution (SS; 100 µg/mL proteinase K for nuclease inactivation, 1% w/v SDS for release of siRNA, 0.5 mM EDTA for protection of RNA from activated proteinase K, and upH_2_O) was added to terminate the reaction and release the siRNA. After incubation with SS at 37°C for 30 min, the samples were loaded onto a 2% agarose gel prepared with 0.5X Tris-Borate-EDTA (TBE) buffer using 50% glycerol as a gel loading reagent. The protected siRNA bands were visualized following ethidium bromide staining using a UV Transilluminator as before. Naked siRNA+SS, cSLN/siRNA complexes treated with SS were used as negative controls to compare with nuclease-treated samples. 50% FBS+SS was loaded as a control to show the bands resulted from serum.

### 2.6. Transmission electron microscopy (TEM)

Samples (3 µL) were dried on carbon-coated 400 mesh copper grids (Electron Microscopy Sciences, USA) overnight at room temperature in a desiccator. TEM imaging was performed in the Transmission Electron Microscopy Laboratory at Middle East Technical University, Ankara, Turkey using high contrast transmission electron microscope Tecnai G2 BioTWIN (FEI Company, USA).

### 2.7. Scanning electron microscopy (SEM)

SEM was performed in The Central Research Laboratory at Izmir Katip Celebi University, Turkey. After drying the samples (5 µL, diluted in upH_2_O) on a glass coverslip overnight at room temperature in a desiccator, they were coated with gold (∼8 nm) under high vacuum using Quorum Q150R ES sputter coater (Quorum Technologies, UK). SEM imaging was carried out at 2 kV by InLens secondary electron detector using Carl ZEISS Sigma 300 VP SEM (ZEISS Group, Germany).

### 2.8. Cell culture

All cell lines were obtained from American Type Culture Collection (ATCC, USA). FBS was heat-inactivated (56°C, 30 min) for cell culture. Prostate cancer cell lines: PC-3 and DU145 were cultured in DMEM/F12 (#31330-038, Gibco) with 5% FBS, 1% penicillin/streptomycin solution (P/S; #15140-122, Gibco); LNCaP was cultured in RPMI-1640 (#R8758, Sigma-Aldrich) with 10% FBS, 1% P/S. All cells were cultured in a humidified atmosphere with 5% CO_2_. Normal prostate epithelial cell lines RWPE-1 and PWR1-E were cultured in K-SFM (#17005-075, Gibco) supplemented with human recombinant epidermal growth factor (2.5 µg), bovine pituitary extract (25 mg), and 1% P/S.

### 2.9. In vitro cellular uptake

#### 2.9.1. Fluorescence microscopy

PC-3 (7 × 10^4^ cells/well) and DU145 (7 × 10^4^ cells/well) cells were seeded into 24-well plates in 1 mL media and cultured until they reached ∼70% confluency prior to treatment. Cells were transfected with 50 nM green fluorescent dye (6-FAM)-labelled siRNA (siGLO; #D-001630-01-05, Dharmacon) in 500 µL media. After 48 h treatment, the nuclei of cells were stained with Hoescht (1 mg/mL) at 1:2000 (v/v) in fresh antibiotic-free medium for 1-2 h at 37°C. Cells were then washed with PBS and images were taken using Lionheart Fx instrument (BioTek, USA).

#### 2.9.2. Flow cytometry

After fluorescence microscopy imaging, cells were trypsinized using standard cell culture techniques and transferred into 5 mL polystyrene round-bottom tubes (#352052, BD Falcon). After PBS-washing and centrifugation (800 rpm, 3 min), the medium was removed, and the cells were re-suspended in 250 µL PBS containing 2% BSA and 1 mM EDTA. A total of 10^4^ cells were counted for each sample and sorted based on green fluorescence positivity using the BD FACS Canto II flow cytometer (BD Biosciences, USA). The data were analyzed by FlowJo, LLC software. The gates were determined for the viable cell population, viable single-cell subpopulation, and FITC (+/−) single-cell subpopulation, respectively. The percentage of green fluorescent cells was normalized to untreated single-cell subpopulation.

### 2.10. siRNA transfection and co-treatment with JIB-04

For EphA2 silencing studies, PC-3 and DU145 cells were seeded into 6-well plates at a density of 1.5 × 10^5^ cells/1.5 mL and 1.75 × 10^5^ cells/1.75 mL, respectively. Cells were cultured until they reached 70% confluency. Three hours before transfection, complete medium was replaced with fresh medium supplemented with FBS. siRNA complexes were prepared at optimal N/P ratios immediately before treatment as described in Section 2.4. Commercial transfection reagent Dharmafect 2 (#T-2002-01, Dharmacon) was used as a positive carrier control according to the manufacturer’s instructions. The complexes of siEphA2 (50 nM; ON-TARGETplus SMARTpool #L-003116-00-0020, Dharmacon) were formed with 5 µL DDAB-cSLN (N/P=10), 4.4 µL DOTMA-cSLN (N/P=8) and 3 µL Dharmafect 2. For other assays, the volumes of the carriers were adjusted by fixing the concentration of siRNA to 50 nM. JIB-04 (260 nM) was dissolved in DMSO (Sigma-Aldrich), gently mixed with antibiotic-free medium (± siRNA complexes), and this mixture was added into the wells. In all experiments; untreated cells (UT), and cells treated with DMSO, empty carriers, or carriers with control siRNA (siControl; ON-TARGETplus Non-targeting Control Pool #D-001810-10-05, Dharmacon) were used as controls.

### 2.11. Determination of mRNA expression levels by quantitative real-time PCR

After 48 h treatment with siEphA2 alone and in combination with JIB-04, total RNA was extracted using the RNeasy Mini Kit (#74104, Qiagen) according to the manufacturer’s instructions including the on-column DNase digestion step (#79254, Qiagen). cDNA was synthesized using a high capacity cDNA reverse transcription kit (Applied Biosystems) using 1 μg total RNA. To determine EphA2 and/or KDM4A mRNA levels, Real-Time PCR was performed on the 7500 Fast Real-Time PCR system (Applied Biosystems) using SYBR Green PCR Master Mix (#4309155, Applied Biosystems). Data were normalized to UT control by using 18S rRNA as a reference housekeeping gene. The results were analyzed using a comparative 2^−ΔΔCt^ method. All primers were purchased from Integrated DNA Technologies (USA). Sequences (5′→3′) of forward (F) and reverse (R) primers were as follows: EphA2_F: GAGTGGCTGGAGTCCATCAA, EphA2_R: TTGAGTCCCAGCAGGCTGTA, KDM4A_F: CCTTGCAAAGCATCACTGCA, KDM4A_R: GGACCACTTCCCCTTCAGCA, 18S_F: GATGGGCGGCGGAAAATAG, 18S_R: GCGTGGATTCTGCATAATGGT.

### 2.12. Determination of protein expression levels by Western blot

PC-3 and DU145 cells were treated and harvested as per Section 2.11. Cell pellet was re-suspended in RIPA Buffer (#9806, CST), incubated for 45 min on ice, and then sonicated at 3 microns amplitude for 20 sec (Soniprep 150, SANYO). Subsequently, samples were centrifuged at 13000*g* for 10 min at 4°C, the supernatant was transferred into new tubes. Protein concentration was determined using a Pierce BCA Assay Kit (ThermoFisher), 30 µg total protein was separated using SDS PAGE, and transferred to a PVDF membrane (#88518, ThermoFisher). Membranes were blocked with 5% non-fat milk in TBS-T (Tris-Buffered-Saline Solution containing 0.1% Tween 20) for 1 h. After overnight incubation with primary antibody (EphA2 mouse anti-human, sc-398832, Santa Cruz) at 1:500 dilution, the membrane was incubated with horseradish peroxidase (HRP)-conjugated anti-mouse secondary antibody (1:12000 dilution) for 1 h at room temperature. HRP-conjugated β-actin (A3854, Sigma-Aldrich) at 1:240000 dilution was used as a loading control. The signal was developed using the enhanced chemiluminescence detection reagent SuperSignal West Pico (#34080, Thermo-Scientific). Images were captured using the Fusion FX (Vilber Lourmat). Densitometric analysis was performed using ImageJ (v1.52i, Wayne Rasband, NIH, USA). Data were normalized to β-actin expression. Full scans of Western blot images are shown in **Fig. S1** and **Fig. S2**.

### 2.13. WST-8 (CCK-8) cell viability assay

#### 2.13.1. Cell viability assay (2D)

PC-3 (1 × 10^4^ cells) and DU145 (1.1 × 10^4^ cells), RWPE-1 and PWR-1E (1.2 × 10^4^ cells) were seeded into flat-bottomed 96-well plates in 100 µL media per well. After the incubation period (40 h for cancer cells and 72 h for normal cells), cells were treated with siEphA2 (50 nM) alone and in combination with JIB-04 (260 nM) in 100 µL fresh antibiotic-free medium for 48 h. At the end of the treatment period, the media was removed and a mixture of 10 µL WST-8 reagent (Dojindo) with 100 µL fresh medium was added to each well. After 4 h incubation at 37°C followed by agitation for 1 min, absorbance was measured at 450 nm and with 650 nm set as a reference wavelength (Versa max microplate reader/ Molecular Devices or EL808 microplate reader/ BioTek). After subtracting the absorbance of blank (only medium), the net absorbance value (A_450_-A_650-_A_blank_) was normalized to UT control value and graphed as percentage cell viability. The half-lethal concentration (LC_50_) value of JIB-04 was calculated by CompuSyn software [33].

#### 2.13.2. Cell viability assay (3D)

The protocol reported by Phung et al. was modified and used for forming spheroids of PC-3 [34]. Briefly, a non-adherent surface was obtained by covering the surface of a U-bottomed 96-well plate with poly(2-hydroxyethyl methacrylate) (PolyHEMA; #P3932, Sigma-Aldrich) dissolved in ethanol (5 mg/mL); and evaporating the ethanol under a non-humidified incubator for 2 days. PC-3 cells were seeded (5 × 10^3^ cells/well) onto the polyHEMA coated plate and centrifuged at 1410 rpm for 10 min immediately. PC-3 spheroids were obtained after incubation for 5 days at 37°C. All steps were followed as stated for 2D cell viability, except treatment time was extended to 96 h instead of 48 h. After adding WST-8 reagent and incubating for 4 h, the mixture of reagent:media was transferred to a flat-bottomed 96-well plate to measure the absorbance. The images of spheroids were recorded using the Lionheart Fx imaging system.

### 2.14. Wound healing assay

PC-3 cells were seeded into a 24-well plate at the density of 1.2 × 10^5^ cells/1 mL and cultured until 100% confluency. A single and straight scratch for each well was formed using 200 µL pipette tips and cells were washed with PBS twice to remove the floating cells. After treatment, the phase-contrast images of the wells were recorded by Lionheart Fx imaging system. Images were taken from the same area of the wells to determine the closure rate of the scratch at 0, 24, 48, and 72 h. The scratch area was calculated using *MRI Wound Healing Tool* plugin of ImageJ software [35].

### 2.15. Clonogenic assay

PC-3 cells were treated in 6-well plates as per Section 2.10 and the clonogenic assay was conducted as previously described [36]. After 48 h treatment, cells were washed with PBS and trypsinized. The cells for each treatment were counted using a hemocytometer after resuspension in a fresh medium. The cells were seeded into new 6-well plates for each treatment at a density of 2500 cells/ 3 mL. These cells were incubated for 10 days without changing the medium. The colonies formed after 10 days were washed with PBS and fixed with ice-cold methanol. Fixed cells were stained with 1 mL 0.5% crystal violet in 25% (v/v) methanol for 20 min. After washing the wells with 4 mL distilled water twice to remove the excess dye, the plates were dried overnight. The colony images at 1200 dpi were recorded by GelCount (Oxford Optronix, UK) instrument. Percentage of colony intensity was calculated using the ColonyArea plugin of ImageJ software [37]. Data were normalized to UT control.

### 2.16. Statistical analysis

Unless otherwise stated, data were representative of at least three independent experiments and values were expressed as mean ± standard error mean (SEM) or mean ± standard deviation (SD). The difference of data between groups was analyzed by ANOVA with a recommended post hoc test by using Graphpad Prism Software-version 6.0 (USA). p values smaller than 0.05 were considered statistically significant.

## 3. Results & Discussion

### 3.1. Characterization of cSLNs and their siRNA complexes

As we aimed passive targeting of nanoparticles to the tumor cells, the particle size should range between 20-200 nm [38]. Besides particle size, narrow size distribution (PDI ≤ 0.5) is required for developing safe, stable, and effective formulation [39]. Here, we successfully prepared two cSLNs with different cationic lipids by modifying the hot microemulsion method, and their cSLN/siRNA complexes via electrostatic interaction (**Fig. 1A**). Both formulation had small particle size (< 200 nm) and narrow size distribution (PDI < 0.4) (**Fig. 1B**). To obtain positively charged nanoparticles, two-tailed cationic lipids DDAB and DOTMA were preferred due to decreased toxicity than their one- or three-tailed counterparts [40,41]. There was an approximately 2-fold increase in particle size of cSLN when cationic lipid DDAB was substituted by DOTMA. Both cationic lipids provided a positive surface charge of more than +30 mV (**Fig. 1B**), which resulted in successful complexation with negatively charged siRNA. As shown in **Fig. 1C**, the optimum N/P ratio was determined as 10 for DDAB-cSLN and 8 for DOTMA-cSLN, and these ratios were used in all subsequent experiments. Although there was a decrease in zeta potential after complexation with siRNA, both complexes preserved their positive charge. Also, particle size and size distribution were maintained in the desired range with siRNA complexes (**Fig. 1B**).

**Fig. 1.**
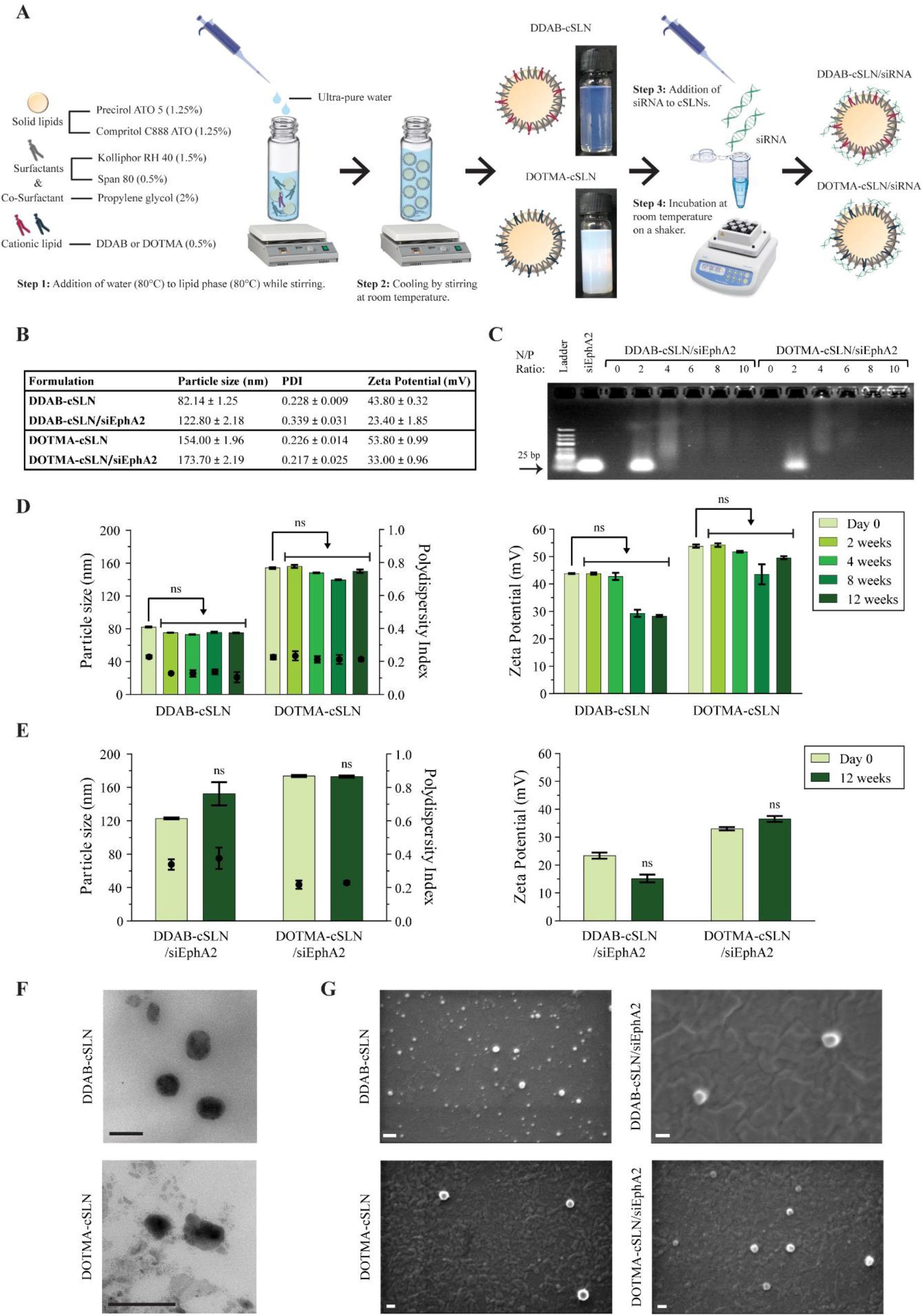
Preparation and characterization of cSLNs and their complexes with siEphA2. (**A**) Overview of steps involved in the preparation of cSLN/siRNA complexes. (**B**) Physicochemical properties of cSLNs and cSLN/siRNA complexes. (**C**) Representative image of gel retardation assay showing complexation of cSLNs with siRNAs at different N/P ratios. Naked siRNA and empty nanoparticles (N/P=0) were used as controls. Time-dependent physicochemical properties of cSLNs (**D**) and cSLN/siRNA complexes (**E**). Bars represent particle size and zeta potential; dots represent PDI. Data are given as mean ± SD of three measurements (ns: not significant, two-way ANOVA followed by Sidak’s test). Representative TEM (**F**) and SEM (**G**) images of cSLNs and cSLN/siRNA complexes, Scale bars, 200 nm.

Owing to their solid lipid matrix, cSLNs are advantageous over other lipid-based carriers such as liposomes and nanoemulsions in terms of storage stability [42]. We evaluated time-dependent changes in the physicochemical properties of cSLNs and complexes. There was no change in size, PDI, and zeta potential values of cSLNs/ complexes after 12-weeks of storage at room temperature (**Fig. 1D & E**). The formation of stable cSLNs against particle agglomeration/aggregation for 12-weeks was probably due to high electrostatic repulsion between cationic particles [43].

Next, we performed TEM and SEM to further characterize the morphology of our formulations. Both nanoparticles displayed a spherical/oval shape (**Fig. 1F & G**) and preserved their morphology even after complexation with siRNA (**Fig. 1G**). These results indicate the spherically-shaped nanoparticles and confirm the results of the DLS measurements.

### 3.2. RNase A and serum stability of cSLN/siRNA complexes

Protecting siRNA from nucleases is one of the prerequisites for an effective nucleic acid delivery system in both *in vitro* and *in vivo* applications. Therefore, we evaluated the siRNA-protection ability of complexes against RNase A and serum nucleases before proceeding to the transfection studies conducted with the serum-containing media to mimic *in vivo* conditions.

As shown in **Fig. 2A**, while naked (free) siRNA was completely degraded in the presence of RNase A, both complexes protected siRNA from complete degradation. Moreover, degradation of naked siRNA started earlier and it was almost completely degraded after 4 h FBS treatment (**Fig. 2B**). However, siRNA was protected from complete degradation by FBS when it was formulated. These data demonstrate that both cSLN formulations decreased the degradation of siRNA by nucleases and support the suitability of the optimum N/P ratios selected for cell culture studies.

**Fig. 2.**
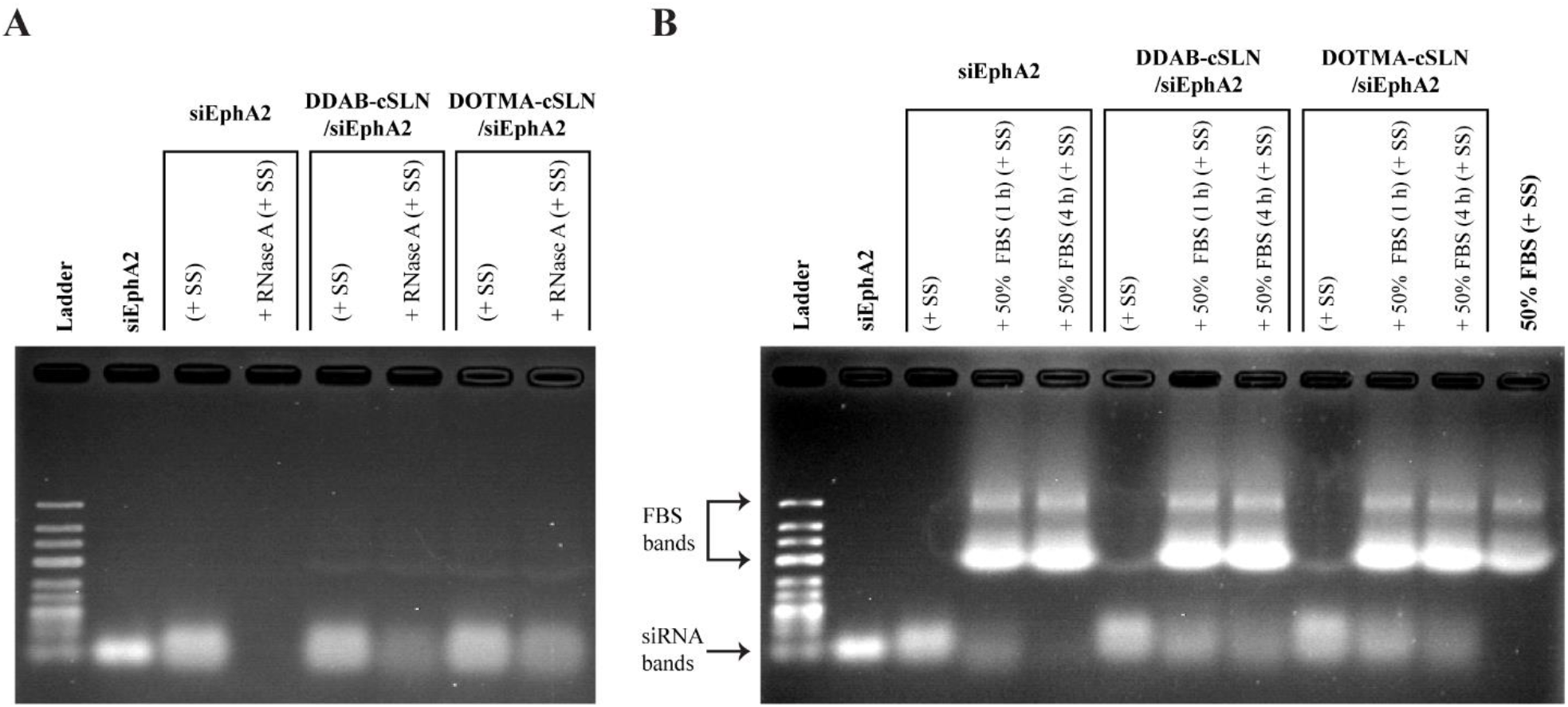
Representative images of agarose gel electrophoresis showing RNase A (**A**) and serum (**B**) stability of cSLN/siEphA2 complexes. cSLN/siRNA complexes were treated with RNase A (30 min) and serum (1 h & 4 h) at 37°C. Electrophoresis was conducted using 2% Agarose gel in 0.5X TBE buffer at 50 V for 70 min. SS: Stop solution. 50% FBS+SS was loaded as a control to show the bands resulted from serum.

### 3.3. In vitro cellular uptake of cSLN/siRNA complexes in DU145 and PC-3 cells

Following characterization and nuclease protection studies, the most effective cSLN formulation for siRNA delivery was determined by cellular uptake efficiency, EphA2 silencing efficiency, and cytotoxicity studies in prostate cancer cell lines. We first measured basal EphA2 expression levels in three commonly used prostate cancer cell lines, LNCaP, DU145, and PC-3. In parallel with the literature [44,45], no EphA2 protein expression was detected in androgen-dependent LNCaP (**Fig. S1**). On the other hand, the basal expression level of the EphA2 was moderate and high in DU145 and PC-3, respectively. The androgen-independent character of these two cell lines has been shown to promote the development of advanced prostate cancer [46]. Therefore, further experiments to compare the efficiency of siEphA2 complexes were conducted in DU145 and PC-3 cell lines.

Next, we performed flow cytometry and fluorescence microscopy analyses to determine the cellular uptake efficiency of cSLN complexes with siGLO. The number of green fluorescent cells detected by flow cytometry shows the number of transfected cells and is expressed as a percentage of cellular uptake efficiency [47]. Moreover, according to the fluorescence microscopy analysis, the homogenous distribution of green fluorescence inside the cell is attributed to the free siGLO that is successfully released from the carrier to the cytosol [47]. Flow cytometry measurements and fluorescence microscopy images showed that Dharmafect 2/siGLO increased the number of green fluorescent cells by 63% in PC-3 (**Fig. 3C**) and 69% in DU145 (**Fig. 3D**), and resulted in a homogenous distribution of green fluorescence in both transfected cells (**Fig. 4**). These data demonstrated that the transfection conditions were optimum in both cell lines.

**Fig. 3.**
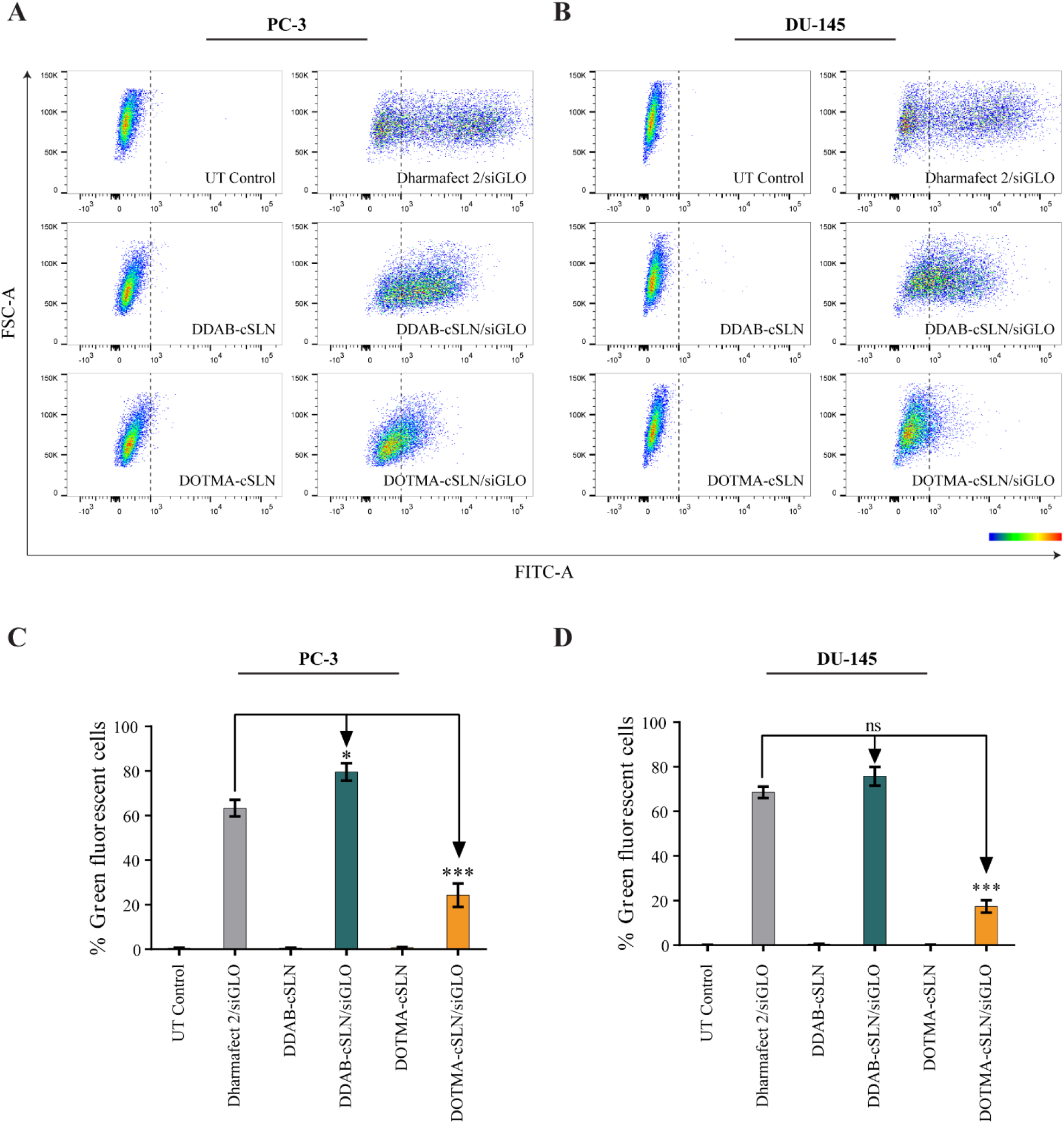
Representative flow cytometry scatter plots (**A & B**) and bar plots (**C & D**) showing cellular uptake efficiency of siGLO complexes in prostate cancer cells (48 h). Color map corresponds to the area of cell density from low (blue) to high (red). (**C & D**) The results are given as the mean± SEM. Asterisks indicate statistical significance (*p<0.05, ***p<0.001, two-way ANOVA followed by Tukey’s test, n = 4).

**Fig. 4.**
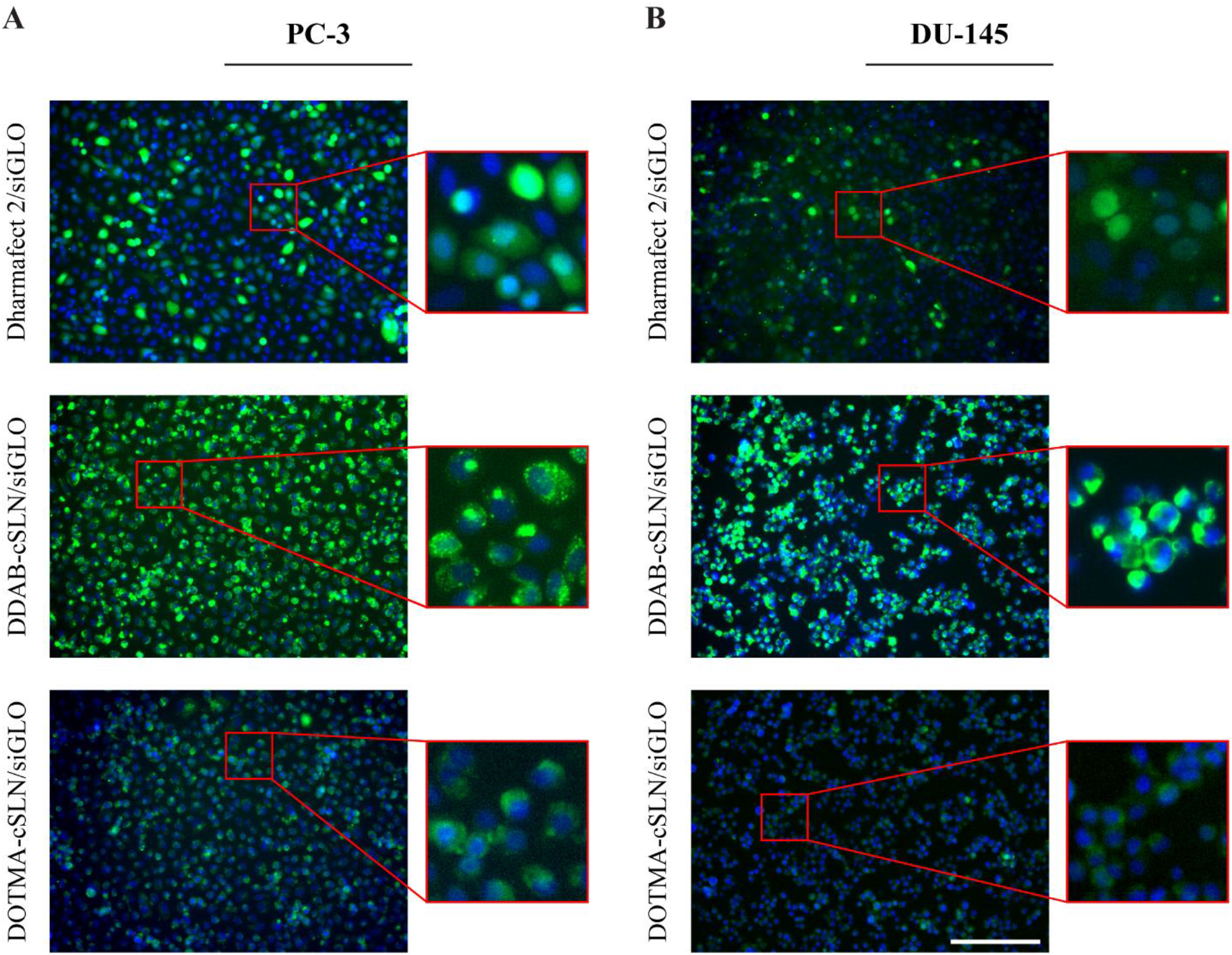
Representative fluorescence microscopy images of PC-3 (**A**) and DU145 (**B**) prostate cancer cells transfected with siGLO complexes (green) and stained with Hoechst dye (blue). Scale bar, 200 µm.

No significant change in green fluorescence levels after treatment of DU145 and PC-3 cells with empty cSLNs indicating that there was no auto-fluorescence effect of these cSLNs (**Fig. 3**). In both cell lines, the highest uptake efficiency was observed with DDAB-cSLN/siGLO complex as compared to other complexes (**Fig. 3C & D**). Notably, its cellular uptake efficiency was even superior to commercially available transfection reagent Dharmafect 2 in PC-3 (**Fig. 3C**). Although the percentage of green fluorescent cells was similar in both cell lines after DDAB-cSLN/siGLO treatment, fluorescence microscopy analysis yielded that the distribution of green fluorescence in the cytoplasm was more homogenous in PC-3 cells (**Fig. 4A**) as compared to DU145 cells (**Fig. 4B**). These data suggest that gene silencing efficiency of DDAB-cSLN might be lower in DU145 cells due to the limited release of siRNA from nanoparticles into the cytosol. Moreover, cellular uptake efficiency of DOTMA-cSLN/siGLO complex was higher in PC-3 (**Fig. 3C**) than in DU145 (**Fig. 3D**). However, it was still significantly lower than other siGLO complexes. Thus, the gene silencing efficiency of DOTMA-cSLN is expected to be relatively low compared to other carriers.

### 3.4. EphA2 silencing efficiency and cytotoxicity of cSLN/siRNA complexes in DU145 and PC-3 cells

To select the optimum cSLN carrier for co-administration studies with JIB-04, gene silencing efficiency and cytotoxicity of siEphA2 complexes were investigated in PC-3 and DU145 cell lines (**Fig. 5**).

**Fig. 5.**
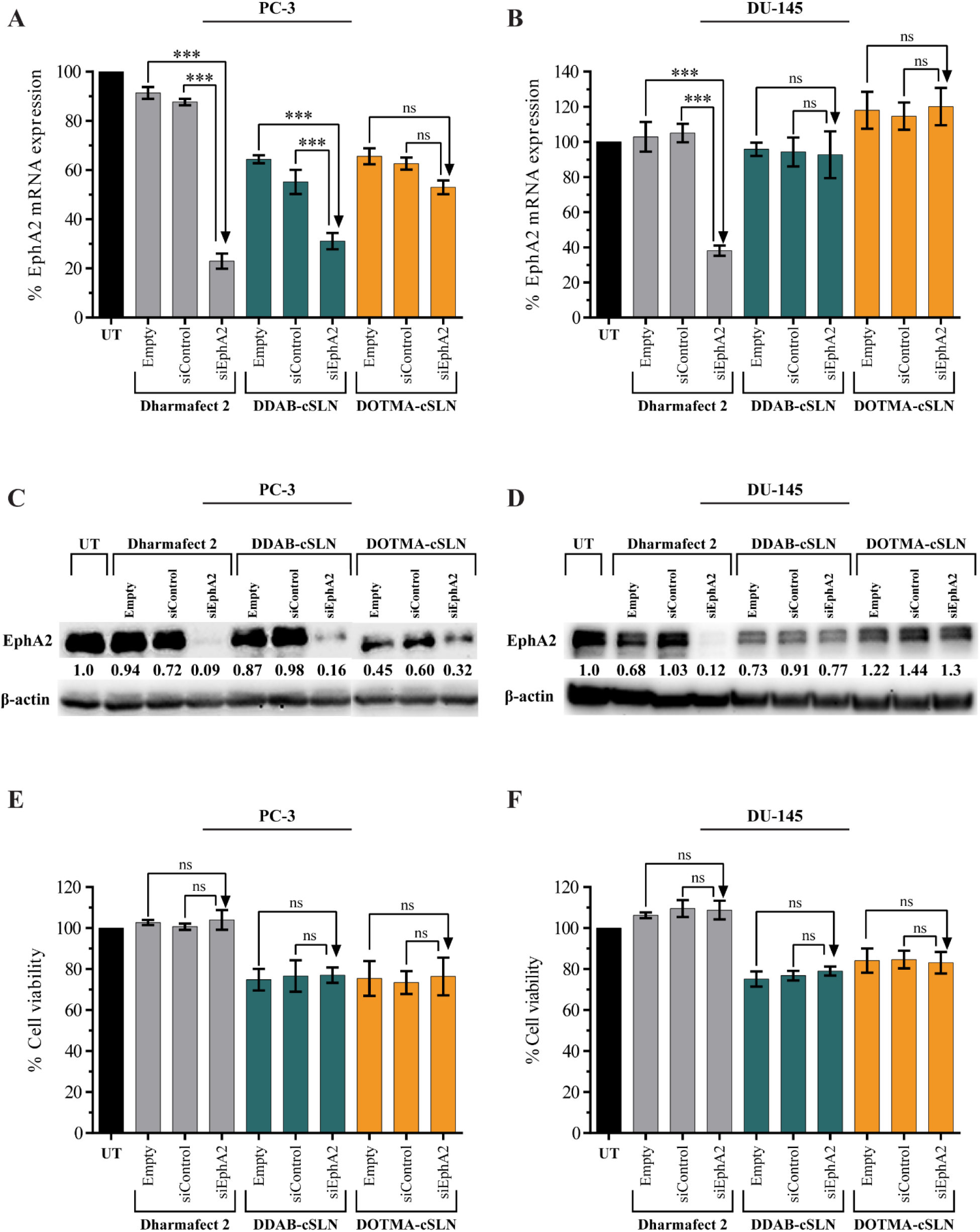
Gene silencing efficiency and cytotoxicity of siEphA2 complexes in DU145 and PC-3 cells (48 h). (**A & B**) Bar plots showing the changes in EphA2 mRNA expression levels determined by qRT-PCR. 18S rRNA was used as a reference gene. (**C & D**) Representative Western blot images and densitometric analysis of EphA2 protein expression levels. β-actin was used as a loading control. (**E & F**) Bar plots showing the effect of cSLNs and their complexes on cell viability. The results are given as mean ± SEM. Asterisks represent statistical significance (UT: Untreated control, ns: not significant, ***p<0.001, one-way ANOVA followed by Tukey’s test, n=3).

According to qRT-PCR measurements (**Fig. 5A & B**), in both cell lines, commercial transfection reagent Dharmafect 2 successfully silenced EphA2 while no significant change was observed with empty carrier and siControl complex. This indicates that 50 nM siRNA concentration was appropriate for silencing studies. Both cSLN/siEphA2 complexes downregulated EphA2 mRNA expression as compared to their empty carrier and non-targeting siControl in PC-3. However, this decrease was only significant with DDAB-cSLN (**Fig. 5A**). Conversely, none of the cSLN/siEphA2 complexes significantly altered the mRNA expression of EphA2 relative to controls in DU145 (**Fig. 5B**). Surprisingly, the administration of empty cSLNs downregulated EphA2 mRNA levels in PC-3, but not in DU145, by an unknown mechanism.

We also verified the EphA2 silencing efficiency by Western blot. The reduction in EphA2 mRNA was accompanied by decreased EphA2 protein expression following 48 h transfection with cSLN/siEphA2 complexes in PC-3 cell lysates (**Fig. 5C**). However, this change in DU145 was not as robust as in PC-3 (**Fig. 5D**). In both cell lines, Dharmafect 2/siEphA2 complex successfully inhibited the EphA2 protein (**Fig. 5C & D**). Since DDAB-cSLN showed similar silencing efficacy to commercial transfection reagent Dharmafect 2 in PC-3, it can be a promising siRNA carrier system for targeting other genes.

Lastly, we investigated the cytotoxicity of carriers and their siRNA complexes applied at the concentrations that gene silencing experiments were performed (**Fig. 5E & F**). While Dharmafect 2 showed no cytotoxic effect, empty cSLNs exhibited modest cytotoxicity on the viability of PC-3 and DU145 cells, which is acceptable for transfection reagents [48]. No significant change was observed in the cytotoxicity of siControl complexes compared to corresponding empty carriers indicating that there was no off-target effect of siRNA on the viability of PC-3 and DU145. Silencing EphA2 showed no significant effect on the viability in both cell lines as compared to siControl treatments.

Although stability against nucleases and toxicity were similar for both cSLNs, we decided to continue the subsequent experiments with the optimum cSLN, DDAB-cSLN, which showed higher cellular uptake and remarkable EphA2 silencing at both mRNA and protein levels in PC-3. Smaller size of DDAB-cSLN might be responsible for its better efficiency than DOTMA-cSLN. However, the well-characterized DOTMA-cSLN may be useful for transfecting other cell lines because the cellular uptake and gene silencing efficiencies of transfection reagents can differ depending on the cell types [49]. To our knowledge, this is the first time cSLN carriers have been developed to deliver siRNA targeting EphA2.

### 3.5. The cytotoxic effect of siEphA2 complexes co-administered with JIB-04 in PC-3 cells

Before proceeding to co-administration studies, the 48 h LC_50_ value of JIB-04 was calculated as 260 nM [*correlation coefficient (r)*=0.9575] in PC-3 cells (**Fig. 6A**). Then, the cytotoxicity of siEphA2 co-administered with JIB-04 was evaluated both in 2D and 3D cell cultures of PC-3 cell line (**Fig. 6B & C**). According to the results from the 2D cell viability assay (**Fig. 6A**), JIB-04 decreased the cell viability by 49%, which was consistent with its calculated LC_50_ value. Next, cytotoxicity of these candidate drugs was also investigated in normal prostate epithelial cell lines RWPE-1 and PWR-1E. JIB-04 did not show any toxicity to normal prostate epithelial cells cancer cells (**Fig. S3**). These findings correspond to similar results in a study performed by L. Wang et al. [23]. When applied under 1 µM concentrations for 96 h, JIB-04 was cytotoxic on prostate cancer cells, however, it was not cytotoxic to normal prostate cells (except SV40 transformed cell line) [23]. When the vehicle control DDAB-cSLN+DMSO was administered to normal prostate epithelial cells, the cell viability was determined as 107% in RWPE-1 (**Fig. S3A**) and 84% in PWR-1E (**Fig. S3B**), whereas it was 77% in PC-3 (**Fig. 6B**). Since higher cytotoxicity was observed in cancer cells than in normal cells, both JIB-04 and DDAB-cSLN formulation can be advantageous as an anticancer drug and an siRNA carrier system, respectively, for prostate cancer therapy.

**Fig. 6.**
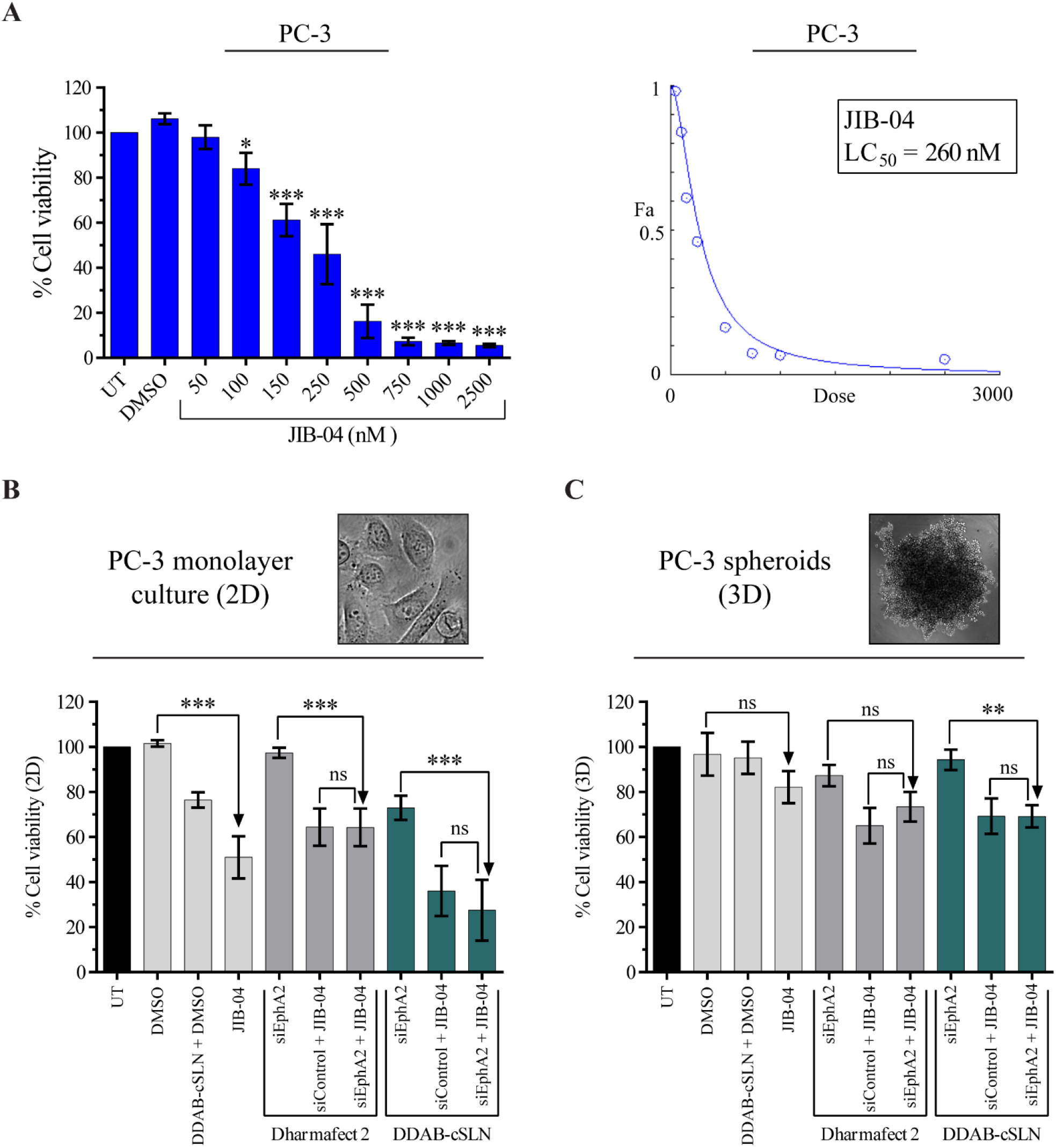
Effect of siEphA2 complexes and JIB-04 on cell viability of PC-3 cells. (**A**) Viability (left) and dose-response curve (right) of PC-3 cells after JIB-04 treatment for 48 h. The results are presented as mean ± SD. Asterisks represent statistical significance (*p<0.05, ***p<0.001, one-way ANOVA followed by Sidak’s test, n=3). The effect of siEphA2 complexes co-administered with JIB-04 on cell viability in PC-3 cells cultured as monolayer cells (2D; 48 h) (**B**) and spheroids (3D; 96 h) (**C**). The results are given as the mean ± SD (UT: Untreated control, ns: not significant, **p<0.01, ***p<0.001, one-way ANOVA followed by Tukey’s test; n=4 for 2D, n=3 for 3D).

In monolayer (2D) PC-3 and DU145 prostate cancer cells (**Fig. 5E & F** and **Fig. 6B**), there was no statistically significant difference in cell viability between siControl and siEphA2 complexes of both carriers. The effect of EphA2 downregulation on cell viability differs depending on the tumor type/subtype. Similar to our results, silencing EphA2 with short hairpin RNA (shRNA) did not cause a significant change in cell viability in salivary adenoid cystic carcinoma (SACC) [50]. On the contrary, cell viability was significantly decreased after siEphA2 treatment in gastric cancer [51], glioma [52], and malignant mesothelioma [53]. Interestingly, silencing siEphA2 decreased the viability of non-metastatic type of renal cell carcinoma (RCC) cells, but this effect was not observed with its metastatic type [54].

When siEphA2 complexes were co-administered with JIB-04 in 2D model of PC-3 cells, no significant change in cell viability was observed compared to siControl+JIB-04 treatments (**Fig. 6B**). This indicates that the effect of JIB-04 on the viability of PC-3 cells was not altered by silencing EphA2. However, cytotoxicity of JIB-04 was increased when combined with siControl or siEphA2 complex of DDAB-cSLN. This effect was probably due to the cytotoxicity of DDAB-cSLN in PC-3.

Tumor spheroids have attracted tremendous attention as they represent a bridge between *in vitro* and *in vivo* toxicity studies [55]. In this study, we also conducted a cytotoxicity assay with PC-3 tumor spheroids cultured three-dimensionally using the non-adherent surface method with polyHEMA. The results of 3D cell viability assay are shown in **Fig. 6C**, and the phase-contrast microscopy images of PC-3 spheroids 48, 72, and 96 h after treatments were provided in **Fig. S4**. Cytotoxicity results obtained in 3D tumor spheroids were mostly parallel to those observed in 2D monolayer cells. Although JIB-04 was applied at its LC_50_ value determined in 2D culture, the viability of PC-3 cells cultured as spheroids was decreased by 18%. Even if this decrease was not statistically significant as in 2D cell viability assay, given that 3D spheroid formation is more physiologically relevant than 2D culture, this tendency might be promising for further studies. As observed with monolayer PC-3 cells, there was no statistically significant difference in cell viability between siControl and siEphA2 complexes of both carriers. Therefore, the consistent decrease in cell viability by co-administration of siEphA2 complexes with JIB-04 compared to administration of siEphA2 complexes alone is attributed to the cytotoxic effect of JIB-04. Besides, DDAB-cSLN+DMSO control was almost non-toxic in 3D culture (the cell viability was 95%). This result suggests that the DDAB-cSLN formulation may be biocompatible *in vivo*.

### 3.6. The effect of siEphA2 complexes co-administered with JIB-04 on migration

Migratory activity is essential for tumor cells to develop metastasis, which causes the spread of the cells from primary tumor to distant parts of the body and the formation of secondary tumors [56]. Thus, we performed a wound healing assay to evaluate the inhibitory effect of siEphA2 complexes and JIB-04 on the migration of PC-3 cells. Scratches were captured at different time points (0, 24, 48, and 72 h), and scratch areas were compared relative to 0 h (**Fig. 7** and **Fig. S5**).

**Fig. 7.**
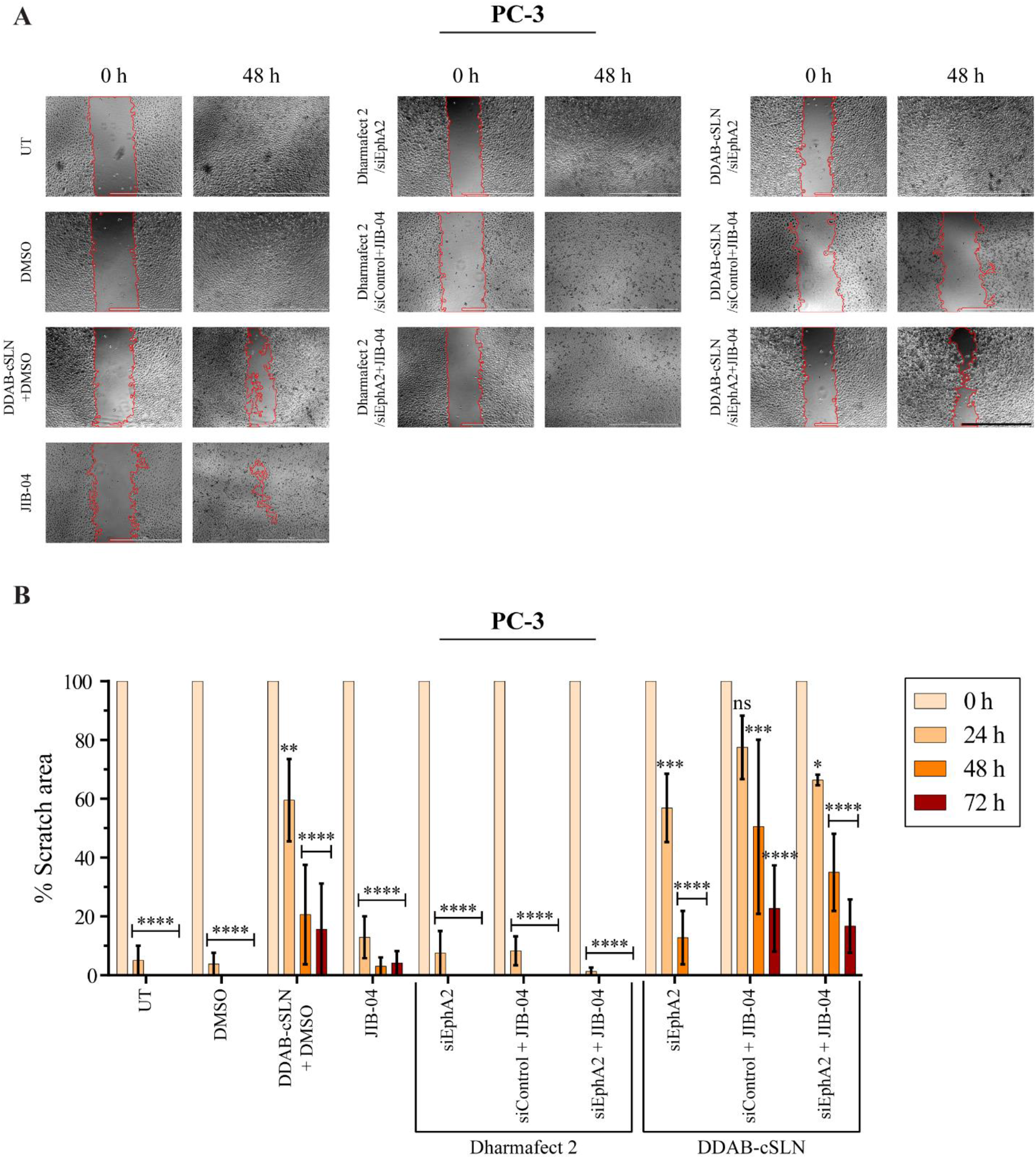
Effect of siEphA2 complexes co-administered with JIB-04 on the migration ability of PC-3 cells. (**A**) Representative phase-contrast images of PC-3 cells showing the changes in the scratch area in response to treatments. *MRI Wound Healing Tool* plugin of ImageJ was used to automatically determine the scratch area. See **Fig. S4** for scratch images at all time points. Scale bar: 1000 µm. (**B**) Bar plots showing the scratch area calculated at 0, 24, 48, and 72 h. Data are shown as mean ± SEM. Asterisks represent statistical significance (UT: Untreated control, ns: not significant, *p<0.05, **p<0.01, ***p<0.001, ****p<0.0001, two-way ANOVA followed by Dunnett’s test; n=3).

As shown in **Fig. 7B**, scratches were almost closed after 24 h in UT and DMSO controls. Treatment with Dharmafect 2/siEphA2 complex did not change this situation. However, the closure of scratch wounds occurred at 48 h with DDAB-cSLN/siEphA2 complex. Therefore, this delay might be due to the DDAB-cSLN carrier, not due to the EphA2 downregulation. This suggestion is also supported by the finding that the scratch was not completely closed after treatment with DDAB-cSLN+DMSO control for 72 h. Even if DDAB-cSLN+DMSO control was less cytotoxic than JIB-04 in PC-3 cells (**Fig. 6B**), DDAB-cSLN formulation led to a greater delay in the migratory activity of PC-3 cells than JIB-04 (**Fig. 7**). Thus, this unexpected migration inhibitory effect of DDAB-cSLN formulation might be independent of its toxicity in PC-3 cells. It may be explained by the ingredients of the DDAB-cSLN formulation inhibiting the migration ability of PC-3 cells. Similar anticancer effect has already been obtained with different excipients in the literature. As demonstrated by Yang et al., D-α-tocopheryl polyethylene glycol succinate (Vitamin E TPGS), which is safely used as an adjuvant in drug formulations, shows a selective anticancer effect by inhibiting multidrug resistance in tumor cells [57]. However, all excipients in DDAB-cSLN formulation should be tested individually and in combinations to investigate this hypothesis, which is outside the scope of this study.

JIB-04 treatment alone delayed the closure time and changed the morphology of the PC-3 cells as compared to UT and DMSO controls after 72 h. Similarly, M. S. Kim et al. demonstrated that JIB-04 decreased the migration and invasion potential of colorectal cancer cells *in vitro* [27]. When JIB-04 was co-administered with DDAB-cSLN/siRNA complexes, the migratory activity of PC-3 cells was lower relative to JIB-04 treatment alone. Again, we attribute this to the migration inhibitory effect of DDAB-cSLN formulation. Moreover, the scratch closed faster with siEphA2 as compared to siControl when complexes of both carriers were co-administered with JIB-04, but this change was not statistically significant. This indicates that silencing EphA2 did not alter the inhibitory effect of JIB-04 on migration ability of PC-3 cells *in vitro*. There are contradictory results in the literature related to *in vitro* migration potential of PC-3 cells following downregulation of EphA2. Similar to our results, Taddei et al. did not observe any alterations in the migration potential of PC-3 cells when they silenced EphA2 by shRNA [58]. However, H. Wang et al. determined a significant decrease in the migration ability of PC-3 cells by using siEphA2 [45]. Although the same cell line was used, these discrepancies may be due to differences in techniques and protocols used in silencing EphA2.

### 3.7. The effect of siEphA2 complexes co-administered with JIB-04 on colony formation

Clonogenic assay is based on testing a single cell for its ability to grow into a colony on a solid surface [36]. This *in vitro* cell survival analysis was performed to investigate the colony formation ability of PC-3 cells after treatment with siEphA2 and JIB-04. Representative images of the colonies and the graph for percentage colony intensity are shown in **Fig. 8**. According to these results, colony intensity was significantly decreased by JIB-04 treatment compared to DMSO control (**Fig. 8B**). This inhibitory effect of JIB-04 on colony formation potential has been reported in other cancer types such as glioblastoma [21,26], lung [25], and Ewing sarcoma [31].

**Fig. 8.**
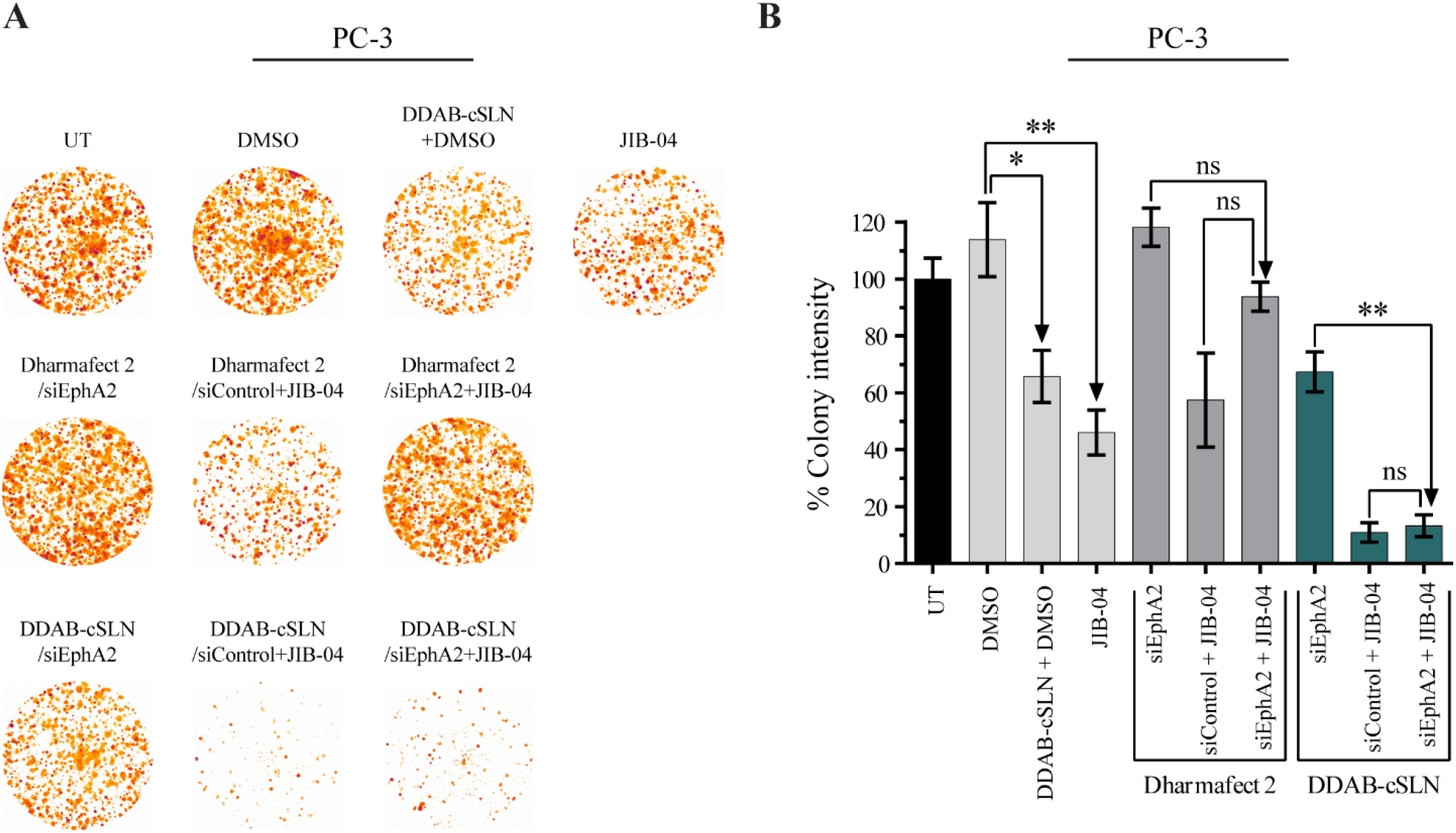
Effect of siEphA2 complexes co-administered with JIB-04 on colony formation ability of PC-3 cells. Representative images (**A**) and bar plots (**B**) of PC-3 colonies after 10 days of incubation. Image processing (thresholding and background subtraction) was done using ImageJ. *Colony Area* plugin of ImageJ was used to automatically quantify the intensity of colonies. The results are given as the mean ± SEM. Asterisks represent statistical significance (UT: Untreated control, ns: not significant, *p<0.05, **p<0.01, one-way ANOVA followed by Tukey’s test, n=3).

Moreover, due to the inhibitory effect of JIB-04 on colony formation, siEphA2+JIB-04 treatment decreased the colony intensity of PC-3 cells as compared to siEphA2 treatment. However, this decrease was only statistically significant when DDAB-cSLN was used as a carrier. Since colony intensity decreased by 34% with only DDAB-cSLN+DMSO treatment, this decrease might result from the cytotoxicity of the formulation in PC-3 cells.

Furthermore, silencing EphA2 caused a tendency of increase in the colony intensity with both carriers (p>0.05). Thus, we considered that silencing EphA2 with siRNA did not alter the colony formation ability of PC-3 cells. To our knowledge, this is the first study showing the effect of siEphA2 alone and in combination with JIB-04 on colony formation ability of prostate cancer cells *in vitro*. In other cancer types such as breast [59], melanoma [60], and Ewing sarcoma [61], the clonogenic growth of cancer cells was suppressed after EphA2 downregulation with siRNA or shRNA. As compared to our results obtained in PC-3 prostate cancer cells, this difference may be due to stable knockdown with shRNA rather than transient silencing with siRNA, and tumor type [62].

### 3.8. The effect of siEphA2 complexes co-administered with JIB-04 on EphA2 and KDM4A expression

We determined the changes in mRNA expression levels of EphA2 and KDM4A in PC-3 cells after co-treatment of siEphA2 complexes with JIB-04. Among the known targets of JIB-04, KDM4A (also known as JMJD2A) was selected due to its link with prostate cancer development and progression. T. D. Kim et al. demonstrated that KDM4A is highly expressed in the metastatic sites of prostate tumors and this overexpression may initiate the development of prostate cancer in mice [63]. Moreover, KDM4A increases the androgen receptor signaling by direct binding [64]. As shown in **Fig. 9A**, the mRNA expression level of KDM4A was significantly decreased by 49% after JIB-04 treatment. However, co-treatment with siEphA2 did not cause a significant change in JIB-04’s inhibitory effect on KDM4A mRNA levels with both carriers (siControl+JIB-04 *vs*. siEphA2+JIB-04).

**Fig. 9.**
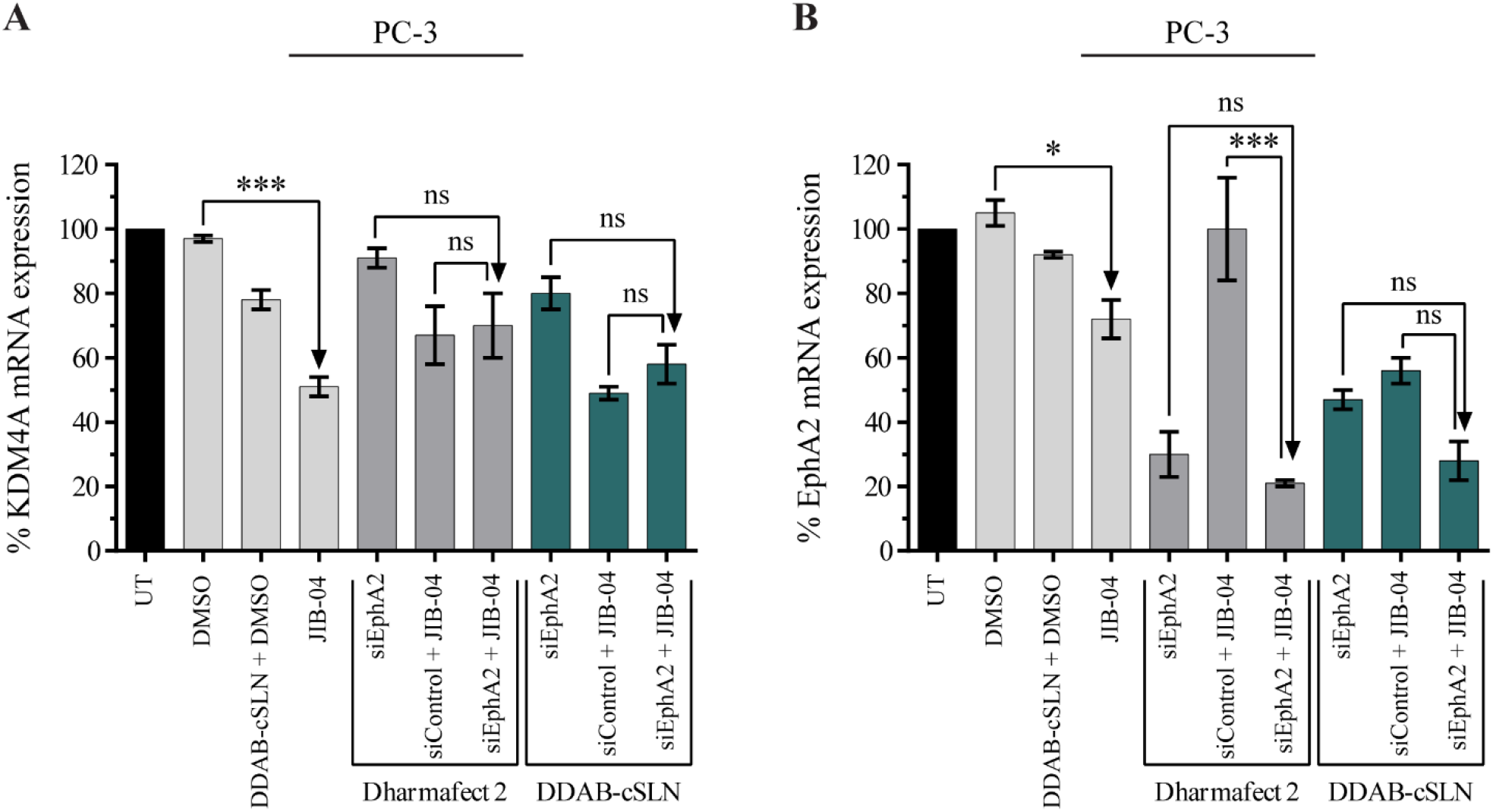
Effect of siEphA2 complexes co-administered with JIB-04 on mRNA expression levels of KDM4A (**A**) and EphA2 (**B**) targets in PC-3 cells (48 h). Bar plots showing the changes in KDM4A and EphA2 mRNA expression levels determined by qRT-PCR. 18S rRNA was used as a reference gene. The results are given as mean ± SEM. Asterisks represent statistical significance (UT: Untreated control, ns: not significant, *p<0.05, ***p<0.001, one-way ANOVA followed by Tukey’s test, n=3).

On the other hand, JIB-04 significantly downregulated EphA2 mRNA expression compared to control, and it also increased the silencing efficiency of siEphA2 complexes (**Fig. 9B**). To our knowledge, this is the first study showing the EphA2 silencing effect of JIB-04, and we suggest that JIB-04 may be a good candidate to combine with the drugs targeting EphA2 to lower the required dose of this drug. However, since silencing EphA2 did not cause any significant change in cell viability, migration, and colony formation abilities of PC-3, further co-administration studies in siEphA2-responsive cancer cells are needed to better investigate the outcomes of EphA2 silencing effect of JIB-04.

Despite several undergoing clinical efforts to develop an effective delivery system targeting EphA2 including prostate cancer, the basic molecular mechanism of Eph signaling is still poorly understood [12]. Thus, there is an urgent need to examine EphA2 response in a well-characterized and highly reproducible *in vitro* models. In this study, we successfully silenced EphA2 using our siRNA delivery system. The results were comparable with Dharmafect 2 as a commercial control. However, we did not observe any change in cellular functions including the viability, migration ability, and clonogenic potential of PC-3 cells after EphA2 silencing. In parallel, it appears that the function of the EphA2 shows paradoxical results highly depending on the context [65]. M. Wang et al. demonstrated that EphA2 may act as a tumor suppressor in SACC after showing that stably silencing EphA2 increased tumor growth, migratory and metastatic potentials both *in vitro* and *in vivo* [50]. Although EphA2 is overexpressed in the skin tumors of both humans and mice, EphA2 knockout in mice increases vulnerability to skin cancer as well as progression to the malignant stage [66]. In contrast, P. Chen et al. demonstrated that overexpression of EphA2 led to an increase in cell viability and invasion in LNCaP prostate cancer cell line stably expressing EphA2. Also, enhanced levels of EphA2 were associated with prostate-specific antigen (PSA) and Gleason score (both are diagnostic/prognostic markers of prostate cancer), lymph node metastasis, and advanced stage of prostate cancer [67]. Furthermore, clinical drug candidate EPHARNA decreased tumor growth by 35-50% after 3-weeks treatment in orthotopic mice models of ovarian cancer [68] and is well-tolerated in mammalian models of mice and Rhesus monkeys [12]. In a study by Duxbury et al., there was no change in cell viability of Capan2 pancreatic adenocarcinoma cells when EphA2 was overexpressed by transient transfection. However, high levels of EphA2 increased Capan2 cells’ invasion potential and resistance to anoikis [69]. Taddei et al. showed that stable EphA2 silencing by shRNA did not affect migration, resistance to anoikis, and adhesion to fibroblast surface in PC-3 cell line. However, anchorage-independent growth was decreased in EphA2-silenced PC-3 by a soft agar colony formation assay. Also, silencing EphA2 reduced the development of bone metastasis in mice [58]. Besides, Miao et al. showed that the migration and invasion ability of PC-3 and glioma cells differed, depending on ligand activation of EphA2 receptor. In their study, EphA2 receptor showed a ligand-dependent tumor suppressor effect by phosphorylating at tyrosine when stimulated via ephrin A1 ligand, while it showed a ligand-independent oncogenic effect when phosphorylated at serine (S897) by Akt [70]. Taken together, in parallel with the literature, our results suggest that tumors would require comprehensive molecular profiling and stratification for effective use of EphA2 signaling as a therapeutic target.

## 4. Conclusion

In summary, we successfully developed a novel DDAB-cSLN/siRNA complex targeting EphA2, which *(i)* has small particle size and size distribution, *(ii)* is biocompatible with normal prostate epithelial cell lines, *(iii)* protects siRNA against nucleases, *(iv)* shows high cellular uptake, *(v)* provides gene silencing as effective as the commercial transfection agent Dharmafect-2 in prostate cancer cell models *in vitro*. Besides, detailed preliminary results of histone lysine demethylase inhibitor JIB-04, alone and in combination with siEphA2-loaded nanoparticles, in prostate cancer cells and tumor spheroids were obtained for the first time. Our siRNA delivery system has a potential use in *in vitro* and *in vivo* studies targeting other genes and cancer types.

## Supporting information

Graphical_Abstract

## Author contributions

E.O. performed the experiments and analyses. All authors (E.O., M.K., A.M.B., S.G.G., B.D.B., E.B., A.G.K., and S.P.F.) made a substantial and intellectual contribution to design the study, and interpretation of data. E.O. wrote the manuscript with critical reviews and editing from all authors. A.G.K. and S.P.F. provided financial support and supervised the study.

### Acknowledgements

Special thanks to Assoc. Prof. PhD Elisabeth Martinez from Department of Pharmacology, University of Texas Southwestern Medical Center for her generous gift, JIB-04. We are grateful to Eamon Breen from the Flow core, Trinity Translational Medicine Institute for assistance with flow cytometry. We thank Department of Pharmaceutical Technology, Ege University for providing access to Zetasizer NanoZS.

## Abbreviations

2D: Two dimensional
3D: Three dimensional
cSLN: Cationic solid lipid nanoparticle
cSLN/siEphA2: Cationic solid lipid nanoparticle loaded with siRNA targeting EphA2 mRNA
DDAB-cSLN: Cationic solid lipid nanoparticle including DDAB
DLS: Dynamic Light Scattering
DOPC: 1,2-dioleoyl-sn-glycero-3-phosphocholine
DOTMA-cSLN: Cationic solid lipid nanoparticle including DOTMA
EphA2: Eph receptor A2
EPR: Enhanced permeability and retention
JIB-04: 5-chloro-N-[(E)-[phenyl(pyridin-2-yl)methylidene]amino]pyridin-2-amine
KDM: Histone lysine demethylase
KRH40: Kolliphor (Cremophor) RH 40
LC_50_: Median lethal concentration which causes the death of 50% of a tested population
N/P ratio: The molar ratio of positively-charged nitrogen (N) groups of cationic lipid to negatively-charged phosphate (P) groups of nucleic acid
PDI: Polydispersity index
PG: Propylene glycol
PolyHEMA: Poly(2-hydroxyethyl methacrylate)
qRT-PCR: Quantitative real-time PCR
RNAi: RNA interference
RTK: Receptor tyrosine kinase
shRNA: Short hairpin RNA
siControl: Negative control siRNA designed to have no known mRNA targets in the cells
siEphA2: siRNA targeting EphA2 mRNA
siGLO: Green fluorescent dye (6-FAM)-labelled siRNA
siRNA: Small interfering RNA
SS: Stop solution for RNase A and serum stability assays
upH_2_O: Ultra-pure water
UT: Untreated control cell population

## References

1. Bray F, Ferlay J, Soerjomataram I, Siegel RL, Torre LA, Jemal A. Global cancer statistics 2018: GLOBOCAN estimates of incidence and mortality worldwide for 36 cancers in 185 countries. CA Cancer J Clin. 2018;68:394–424.

2. Nurgali K, Jagoe RT, Abalo R. Editorial: Adverse effects of cancer chemotherapy: Anything new to improve tolerance and reduce sequelae? Front Pharmacol. 2018;9:245.

3. Ahuja N, Easwaran H, Baylin SB, Ahuja N, Easwaran H, Baylin SB. Harnessing the potential of epigenetic therapy to target solid tumors Find the latest version?: Review series Harnessing the potential of epigenetic therapy to target solid tumors. J Clin Invest. 2014;124:56–63.

4. Mansoori B, Shotorbani SS, Baradaran B. RNA interference and its role in cancer therapy. Adv Pharm Bull. 2014;4:313–21.

5. Chen X, Mangala LS, Rodriguez-Aguayo C, Kong X, Lopez-Berestein G, Sood AK. RNA interference-based therapy and its delivery systems. Cancer Metastasis Rev. Cancer and Metastasis Reviews; 2018;37:107–24.

6. Qi Y, Wang D, Wang D, Jin T, Yang L, Wu H, et al. HEDD: The human epigenetic drug database. Database. 2016;2016:1–10.

7. Setten RL, Rossi JJ, Han Sping. The current state and future directions of RNAi-based therapeutics. Nat Rev Drug Discov. 2019;18:421–46.

8. Kim B, Park JH, Sailor MJ. Rekindling RNAi Therapy: Materials Design Requirements for In Vivo siRNA Delivery. Adv Mater. 2019;31:1–23.

9. Rüger J, Ioannou S, Castanotto D, Stein CA. Oligonucleotides to the (Gene) Rescue: FDA Approvals 2017–2019. Trends Pharmacol Sci. 2020;41:27–41.

10. Parashar D, Rajendran V, Shukla R, Sistla R. Lipid-based nanocarriers for delivery of small interfering RNA for therapeutic use. Eur J Pharm Sci. 2020;142:105159.

11. Subhan MA, Torchilin VP. Efficient nanocarriers of siRNA therapeutics for cancer treatment. Transl Res. 2019;214:62–91. Available from: https://doi.org/10.1016/j.trsl.2019.07.006

12. Wagner MJ, Mitra R, Mcarthur MJ, Baze W, Barnhart K, Wu SY, et al. Preclinical mammalian safety studies of EPHARNA (DOPC Nanoliposomal EphA2-Targeted siRNA). Mol Cancer Ther. 2017;16:1114–23.

13. Yingchoncharoen P, Kalinowski DS, Richardson DR. Lipid-based drug delivery systems in cancer therapy: What is available and what is yet to come. Pharmacol Rev. 2016;68:701–87.

14. Kinch MS, Carles-Kinch K. Overexpression and functional alterations of the EphA2 tyrosine kinase in cancer. Clin Exp Metastasis. 2003;20:59–68.

15. Tandon M, Vemula V, Mittal S. Emerging strategies for EphA2 receptor targeting for cancer therapeutics. Expert Opn Ther Targets. 2011.

16. Ozcan G, Ozpolat B, Coleman RL, Sood AK, Lopez-Berestein G. Preclinical and clinical development of siRNA-based therapeutics. Adv. Drug Deliv. Rev. 2015;87:108–19. Available from: http://www.ncbi.nlm.nih.gov/pubmed/25666164

17. Sharma S V., Lee DY, Li B, Quinlan MP, Takahashi F, Maheswaran S, et al. A Chromatin-Mediated Reversible Drug-Tolerant State in Cancer Cell Subpopulations. Cell. 2010;141:69–80.

18. Berry WL, Janknecht R. KDM4/JMJD2 Histone demethylases: Epigenetic regulators in cancer cells. Cancer Res. 2013;73:2936–42.

19. Lee DH, Kim GW, Jeon YH, Yoo J, Lee SW, Kwon SH. Advances in histone demethylase KDM4 as cancer therapeutic targets. FASEB J. 2020;00:1–24.

20. McAllister TE, England KS, Hopkinson RJ, Brennan PE, Kawamura A, Schofield CJ. Recent Progress in Histone Demethylase Inhibitors. J Med Chem. 2016;59:1308–29.

21. Romani M, Daga A, Forlani A, Pistillo MP, Banelli B. Targeting of Histone Demethylases KDM5A and KDM6B Inhibits the Proliferation of Temozolomide-Resistant Glioblastoma Cells. Cancers (Basel). 2019;11:1–16.

22. Dalvi MP, Wang L, Zhong R, Kollipara RK, Park H, Bayo J, et al. Taxane-Platin-Resistant Lung Cancers Co-develop Hypersensitivity to JumonjiC Demethylase Inhibitors. Cell Rep. 2017;19:1669–84. Available from: http://dx.doi.org/10.1016/j.celrep.2017.04.077

23. Wang L, Chang J, Varghese D, Dellinger M, Kumar S, Best AM, et al. A small molecule modulates Jumonji histone demethylase activity and selectively inhibits cancer growth. Nat Commun. 2013;4:2035. Available from: http://www.nature.com/doifinder/10.1038/ncomms3035

24. Plch J, Hrabeta J, Eckschlager T. KDM5 demethylases and their role in cancer cell chemoresistance. Int J Cancer. 2019;144:221–31.

25. Bayo J, Tran TA, Wang L, Peña-Llopis S, Das AK, Martinez ED. Jumonji Inhibitors Overcome Radioresistance in Cancer through Changes in H3K4 Methylation at Double-Strand Breaks. Cell Rep. 2018;25:1040–1050.e5.

26. Banelli B, Daga A, Forlani A, Allemanni G, Marubbi D, Pistillo MP, et al. Small molecules targeting histone demethylase genes (KDMs) inhibit growth of temozolomide-resistant glioblastoma cells. Oncotarget. 2017;8:34896–910.

27. Kim MS, Cho HI, Yoon HJ, Ahn Y-H, Park EJ, Jin YH, et al. JIB-04, A Small Molecule Histone Demethylase Inhibitor, Selectively Targets Colorectal Cancer Stem Cells by Inhibiting the Wnt/β-Catenin Signaling Pathway. Sci Rep. 2018;8:6611. Available from: http://www.ncbi.nlm.nih.gov/pubmed/29700375

28. Xu W, Zhou B, Zhao X, Zhu L, Xu J, Jiang Z, et al. KDM5B demethylates H3K4 to recruit XRCC1 and promote chemoresistance. Int J Biol Sci. 2018;14:1122–32.

29. Bayo J, Fiore EJ, Dominguez LM, Real A, Malvicini M, Rizzo M, et al. A comprehensive study of epigenetic alterations in hepatocellular carcinoma identifies potential therapeutic targets. J Hepatol. 2019;71:78–90. Available from: https://doi.org/10.1016/j.jhep.2019.03.007

30. Mar BG, Chu SH, Kahn JD, Krivtsov A V., Koche R, Castellano CA, et al. SETD2 alterations impair DNA damage recognition and lead to resistance to chemotherapy in leukemia. Blood. 2017;130:2631–41.

31. Parrish JK, McCann TS, Sechler M, Sobral LM, Ren W, Jones KL, et al. The Jumonji-domain histone demethylase inhibitor JIB-04 deregulates oncogenic programs and increases DNA damage in Ewing Sarcoma, resulting in impaired cell proliferation and survival, and reduced tumor growth. Oncotarget. 2018;9:33110–23.

32. Oner E, Kotmakci M, Kantarci AG. A promising approach to develop nanostructured lipid carriers from solid lipid nanoparticles: preparation, characterization, cytotoxicity and nucleic acid binding ability. Pharm Dev Technol. 2020;25:936–48. Available from: https://www.tandfonline.com/doi/full/10.1080/10837450.2020.1759630

33. Chou T-C, Martin N. CompuSyn for Drug Combinations: PC Software and User’s Guide: A Computer Program for Quantitation of Synergism and Antagonism in Drug Combinations, and the Determination of IC50 and ED50 and LD50 Values. 2005. Available from: http://www.combosyn.com/index.html

34. 1. Phung YT, Barbone D, Broaddus VC, Ho M. Rapid generation of in vitro multicellular spheroids for the study of monoclonal antibody therapy. J Cancer. 2011;2:507–14.

35. Venter C, Niesler CU. Rapid quantification of cellular proliferation and migration using ImageJ. Biotechniques. 2019;66:99–102.

36. Franken NAP, Rodermond HM, Stap J, Haveman J, Bree C Van. Clonogenic assay of cells in vitro. 2006;1:2315–9.

37. Guzmán C, Bagga M, Kaur A, Westermarck J, Abankwa D. ColonyArea: An ImageJ Plugin to Automatically Quantify Colony Formation in Clonogenic Assays. PLoS One. 2014;9:1–9. Available from: https://dx.plos.org/10.1371/journal.pone.0092444

38. Kanasty R, Dorkin JR, Vegas A, Anderson D. Delivery materials for siRNA therapeutics. Nat Mater. 2013;12:967–77.

39. Kaur IP, Bhandari R, Bhandari S, Kakkar V. Potential of solid lipid nanoparticles in brain targeting. J Control Release. 2008;127:97–109.

40. Zhi D, Zhang S, Wang B, Zhao Y, Yang B, Yu S. Transfection efficiency of cationic lipids with different hydrophobic domains in gene delivery. Bioconjug Chem. 2010;21:563–77.

41. Tabatt K, Sameti M, Olbrich C, Müller RH, Lehr CM. Effect of cationic lipid and matrix lipid composition on solid lipid nanoparticle-mediated gene transfer. Eur J Pharm Biopharm. 2004;57:155–62.

42. Teixeira MC, Carbone C, Souto EB. Beyond liposomes: Recent advances on lipid based nanostructures for poorly soluble/poorly permeable drug delivery. Prog Lipid Res. 2017;68:1–11. Available from: http://dx.doi.org/10.1016/j.plipres.2017.07.001

43. Buck J, Grossen P, Cullis PR, Huwyler J, Witzigmann D. Lipid-based DNA therapeutics: Hallmarks of non-viral gene delivery. ACS Nano. 2019;13:3754–82.

44. Walker-Daniels J, Coffman K, Azimi M, Rhim JS, Bostwick DG, Snyder P, et al. Overexpression of the EphA2 tyrosine kinase in prostate cancer. Prostate. 1999;41:275–80.

45. Wang H, Lin H, Pan J, Mo C, Zhang F, Huang B, et al. Vasculogenic mimicry in Prostate cancer: The roles of EphA2 and PI3K. J Cancer. 2016;7:1114–24.

46. Sahin I, Mega AE, Carneiro BA. Androgen receptor-independent prostate cancer: an emerging clinical entity. Cancer Biol Ther. 2018;19:347–8. Available from: https://doi.org/10.1080/15384047.2018.1423926

47. Breunig M, Hozsa C, Lungwitz U, Watanabe K, Umeda I, Kato H, et al. Mechanistic investigation of poly(ethylene imine)-based siRNA delivery: Disulfide bonds boost intracellular release of the cargo. J Control Release. 2008;130:57–63.

48. Wang T, Larcher LM, Ma L, Veedu RN. Systematic screening of commonly used commercial transfection reagents towards efficient transfection of single-stranded oligonucleotides. Molecules. 2018;23:2564.

49. Neuhaus B, Tosun B, Rotan O, Frede A, Westendorf AM, Epple M. Nanoparticles as transfection agents: A comprehensive study with ten different cell lines. RSC Adv. Royal Society of Chemistry; 2016;6:18102–12.

50. Wang M, Zhao X-P, Xu Z, Yan T-L, Song Y, Song K, et al. EphA2 silencing promotes growth, migration, and metastasis in salivary adenoid cystic carcinoma: in vitro and in vivo study. Am J Transl Res. 2016;8:1518–29. Available from: http://www.ncbi.nlm.nih.gov/pubmed/27186278

51. Yuan W, Chen Z, Chen Z, Wu S, Guo J, Ge J, et al. Silencing of EphA2 inhibits invasion of human gastric cancer SGC-7901 cells in vitro and in vivo. Neoplasma. 2012;59:105–13. Available from: http://www.elis.sk/index.php?page=shop.product_details&flypage=flypage.tpl&product_id=2534&category_id=85&option=com_virtuemart

52. Zhou Z, Yuan X, Li Z, Tu H, Li D, Qing J, et al. RNA interference targeting EphA2 inhibits proliferation, induces apoptosis, and cooperates with cytotoxic drugs in human glioma cells. Surg Neurol. 2008;70:562–8. Available from: http://dx.doi.org/10.1016/j.surneu.2008.04.031

53. Nasreen N, Mohammed KA, Antony VB. Silencing the receptor EphA2 suppresses the growth and haptotaxis of malignant mesothelioma cells. Cancer. 2006;107:2425–35.

54. Cho MC, Cho SY, Yoon CY, Lee SB, Kwak C, Kim HH, et al. EphA2 is a potential player of malignant cellular behavior in non-metastatic renal cell carcinoma cells but not in metastatic renal cell carcinoma cells. PLoS One. 2015;10:1–13.

55. Langhans SA. Three-dimensional in vitro cell culture models in drug discovery and drug repositioning. Front Pharmacol. 2018;9:1–14.

56. Palm D, Lang K, Brandt B, Zaenker KS, Entschladen F. In vitro and in vivo imaging of cell migration: Two interdepending methods to unravel metastasis formation. Semin Cancer Biol. 2005;15:396–404.

57. Yang C, Wu T, Qi Y, Zhang Z. Recent advances in the application of vitamin E TPGS for drug delivery. Theranostics. 2018;8:464–85.

58. Taddei ML, Parri M, Angelucci A, Bianchini F, Marconi C, Giannoni E, et al. EphA2 induces metastatic growth regulating amoeboid motility and clonogenic potential in prostate carcinoma cells. Mol Cancer Res. 2011;9:149–60.

59. Lévêque R, Corbet C, Aubert L, Guilbert M, Lagadec C, Adriaenssens E, et al. ProNGF increases breast tumor aggressiveness through functional association of TrkA with EphA2. Cancer Lett. 2019;449:196–206. Available from: https://doi.org/10.1016/j.canlet.2019.02.019

60. Udayakumar D, Zhang G, Ji Z, Njauw CN, Mroz P, Tsao H. Epha2 is a critical oncogene in melanoma. Oncogene. 2011;30:4921–9.

61. Garcia-Monclús S, López-Alemany R, Almacellas-Rabaiget O, Herrero-Martín D, Huertas-Martinez J, Lagares-Tena L, et al. EphA2 receptor is a key player in the metastatic onset of Ewing sarcoma. Int J Cancer. 2018;143:1188–201.

62. Moore CB, Guthrie EH, Huang MTH, Taxman DJ. Short hairpin RNA (shRNA): design, delivery, and assessment of gene knockdown. Methods Mol Biol. 2010;629:141–58.

63. Kim TD, Jin F, Shin S, Oh S, Lightfoot SA, Grande JP, et al. Histone demethylase JMJD2A drives prostate tumorigenesis through transcription factor ETV1. J Clin Invest. 2016;126:706–20.

64. Shin S, Janknecht R. Activation of androgen receptor by histone demethylases JMJD2A and JMJD2D. Biochem Biophys Res Commun. 2007;359:742–6.

65. Pasquale EB. Eph receptors and ephrins in cancer: Bidirectional signalling and beyond. Nat Rev Cancer. 2010;10:165–80.

66. Guo H, Miao H, Gerber L, Singh J, Denning MF, Gilliam AC, et al. Disruption of EphA2 receptor tyrosine kinase leads to increased susceptibility to carcinogenesis in mouse skin. Cancer Res. 2006;66:7050–8.

67. Chen P, Huang Y, Zhang B, Wang Q, Bai P. EphA2 enhances the proliferation and invasion ability of LNCaP prostate cancer cells. Oncol Lett. 2014;8:41–6. Available from: http://www.ncbi.nlm.nih.gov/pubmed/24959216

68. Landen Jr CN, Chavez-Reyes A, Bucana C, Schmandt R, Deavers MT, Lopez-Berestein G, et al. Therapeutic EphA2 Gene Targeting In vivo Using Neutral. Cancer Res. 2005;65:6910–8. Available from: http://cancerres.aacrjournals.org/content/65/15/6910.full.pdf+html

69. Duxbury MS, Ito H, Zinner MJ, Ashley SW, Whang EE. EphA2: A determinant of malignant cellular behavior and a potential therapeutic target in pancreatic adenocarcinoma. Oncogene. 2004;23:1448–56.

70. Miao H, Li DQ, Mukherjee A, Guo H, Petty A, Cutter J, et al. EphA2 Mediates Ligand-Dependent Inhibition and Ligand-Independent Promotion of Cell Migration and Invasion via a Reciprocal Regulatory Loop with Akt. Cancer Cell. 2009;16:9–20. Available from: http://dx.doi.org/10.1016/j.ccr.2009.04.009

