## Supplemental_file for "Development of EphA2 siRNA-loaded lipid nanoparticles and combination with a small-molecule histone demethylase inhibitor in prostate cancer cells and tumor spheroids"

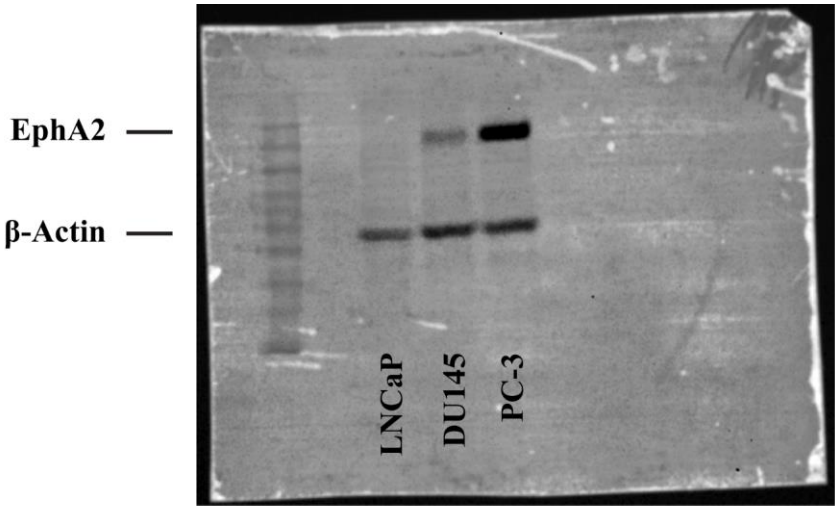


**Fig. S1.** Representative Western blot image showing the basal expression levels of EphA2 protein expression in LNCaP, DU145, and PC-3 prostate cancer cells.
β-actin was used as a loading control. Each well was loaded with 50 μg of total protein. Marker bands: 175, 130, 95, 70, 62, 51, 42, 29, 22, 14 kDa.

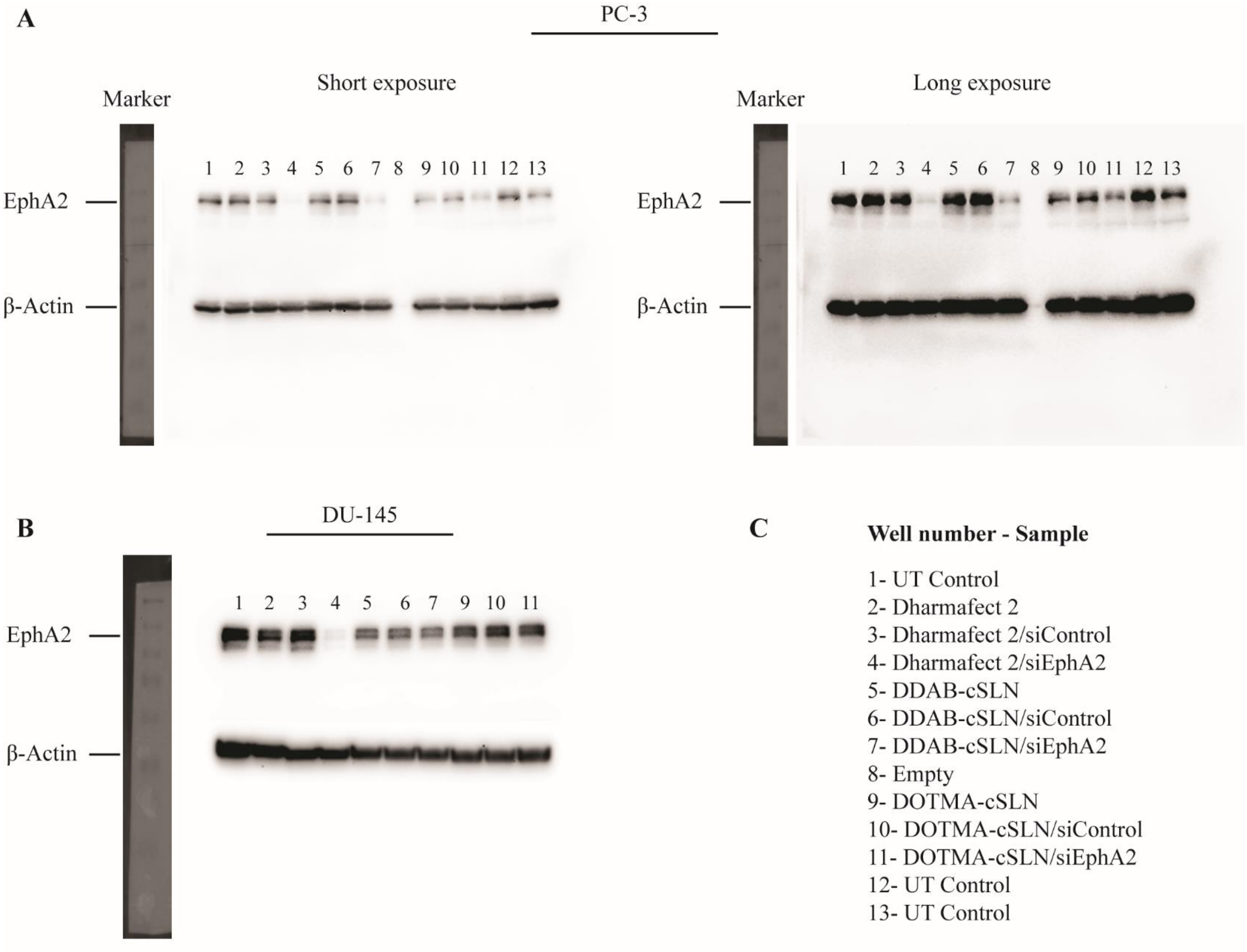


**Fig. S2.** Full scans of Western blot images corresponding to Fig 5C & D. Marker bands: 250, 150, 100 75, 50, 37, 25, 20 kDa.

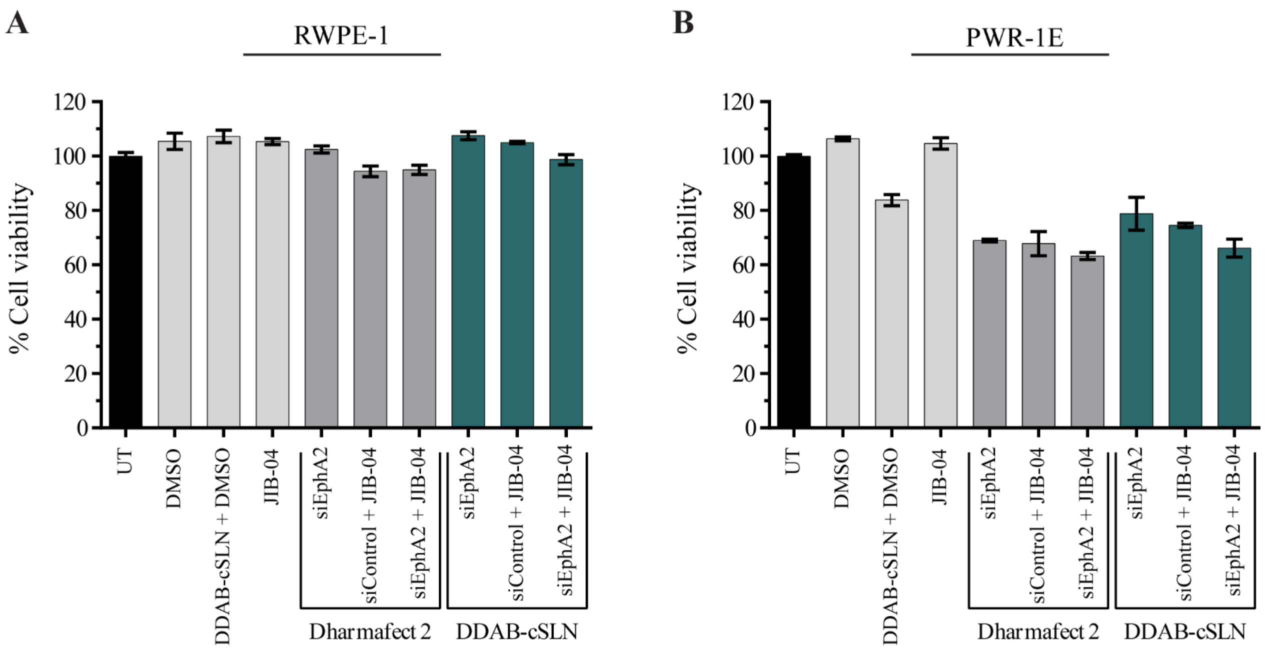


**Fig. S3.** The effect of siEphA2 complexes co-administered with JIB-04 on cell viability in RWPE-1 (**A**) and PWR-1E (**B**) normal human prostate epithelial cell lines (48 h). Data are given as mean ± SD of three measurements.

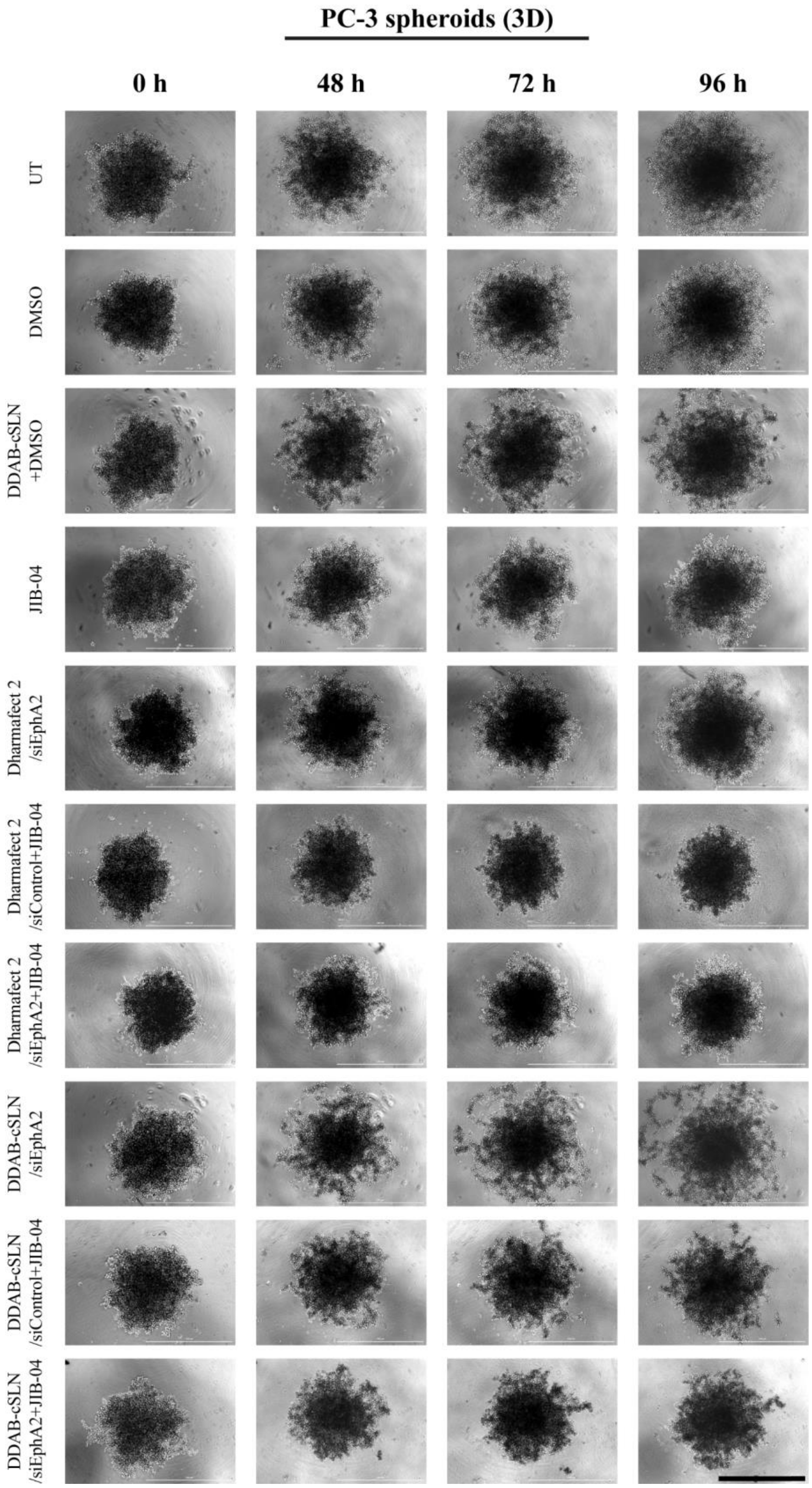


**Fig. S4.** The morphological changes in PC-3 spheroids after co-treatment of siEphA2 complexes with JIB-04. Scale bar, 1000 μm.

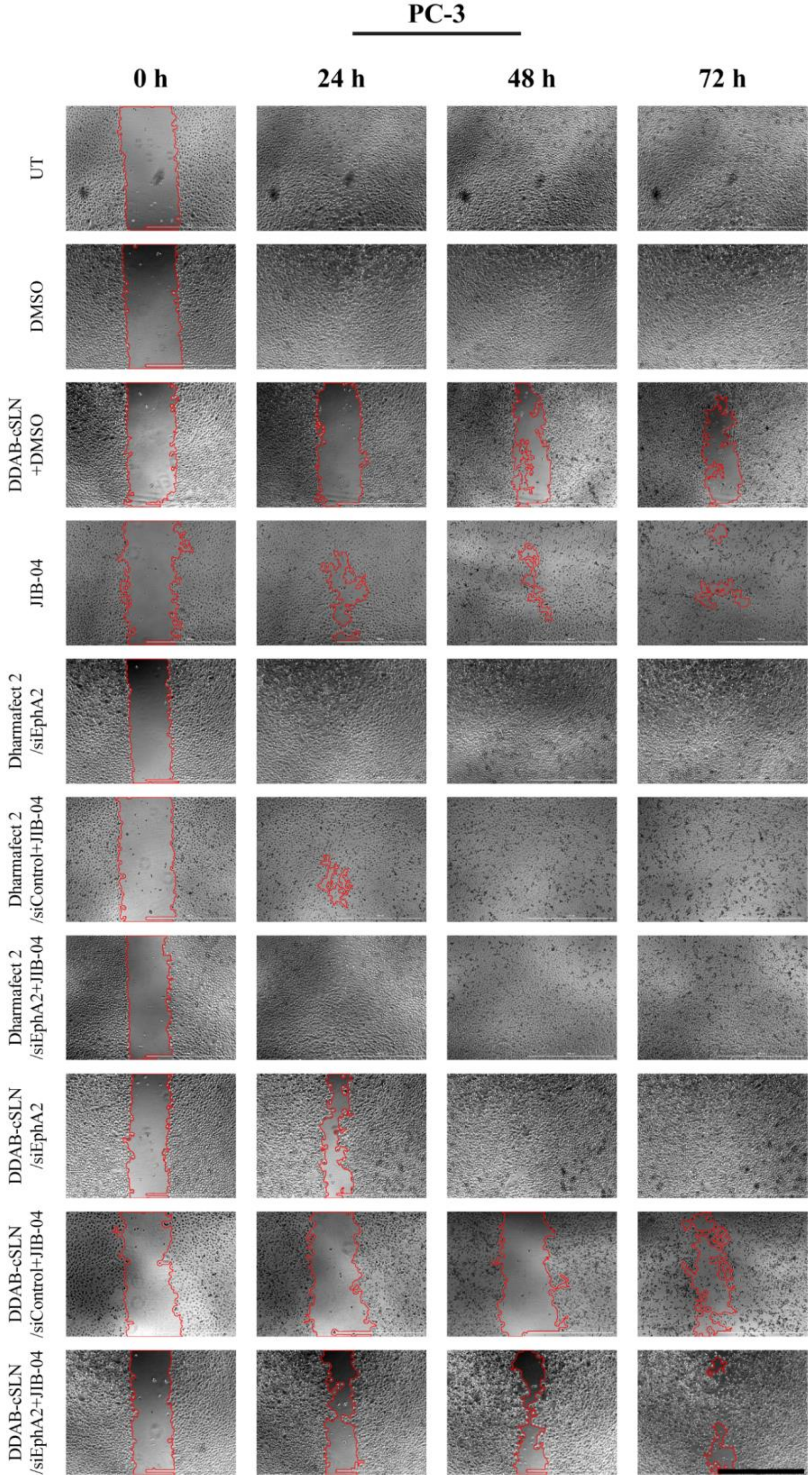


**Fig. S5.** Representative phase-contrast images of PC-3 cells showing the changes in scratch area in response to treatments after 0 h, 24 h, 48 h, and 72 h. *MRI Wound Healing Tool* plugin of ImageJ was used to automatically draw the scratch area. Scale bar, 1000 µm.
