## Supplementary figures and images for "Development of EphA2 siRNA-loaded lipid nanoparticles and combination with a small-molecule histone demethylase inhibitor in prostate cancer cells and tumor spheroids"

### Graphical_Abstract

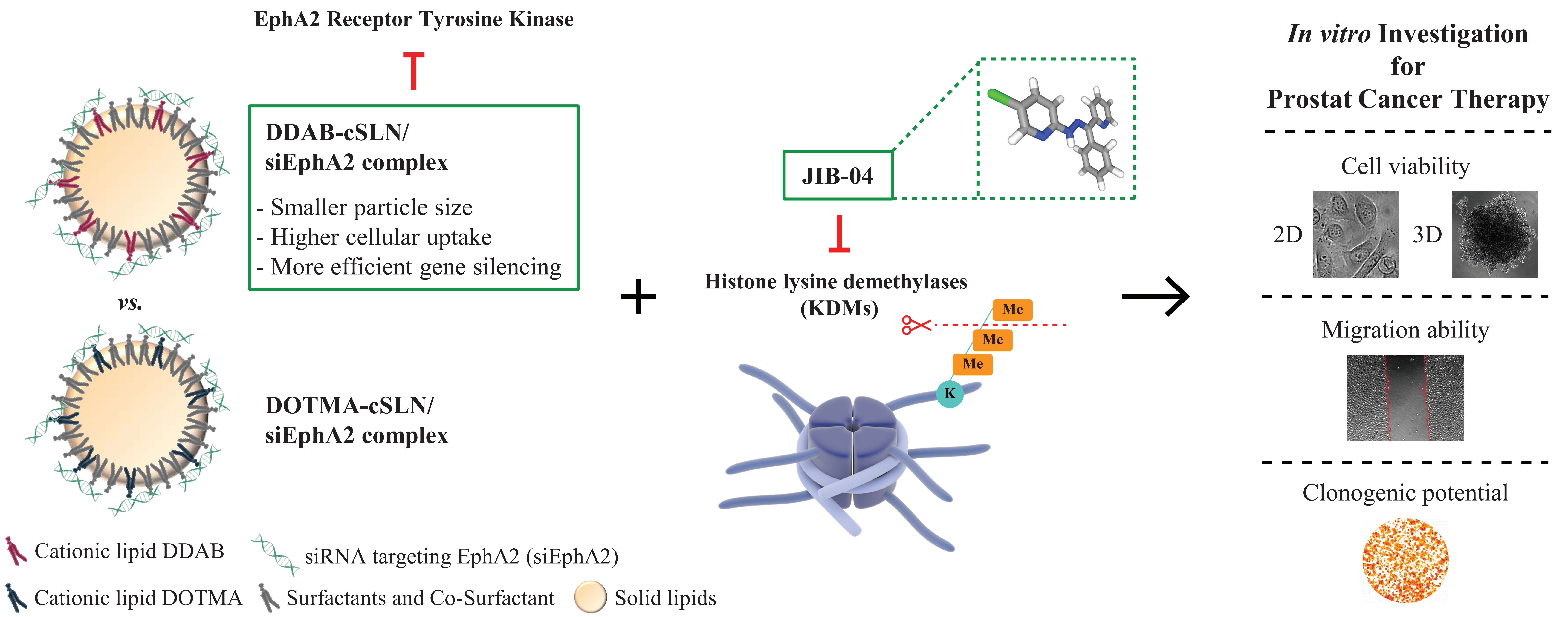
